# NUMonomer enables accurate and scalable nucleic acid structure prediction from primary sequence alone

**DOI:** 10.64898/2026.07.20.739453

**Authors:** Yunda Si, Suqi Zhang, Luonan Chen

**Affiliations:** Key Laboratory of Systems Health Science of Zhejiang Province, School of Life Science, Hangzhou Institute for Advanced Study, University of Chinese Academy of Sciences, Hangzhou 310024, China; College of Electrical Engineering, Naval University of Engineering, Wuhan 430033, China; School of Mathematical Sciences and School of AI, Shanghai Jiao Tong University, Shanghai 200240, China

## Abstract

Accurate and efficient prediction of three-dimensional nucleic acid structures can accelerate functional characterization and enable downstream applications. Recent deep-learning methods have substantially improved nucleic acid structure prediction by incorporating auxiliary inputs such as multiple sequence alignments, secondary-structure annotations, and representations from pretrained language models. However, prediction accuracy remains limited, and generating these auxiliary inputs can be computationally expensive. Here we show that learning the hierarchical organization of experimentally determined structures across multiple scales, from recurring local conformations to global fold topologies, together with exploiting representations shared between RNA and single-stranded DNA, improves model generalization. Guided by these findings, we developed NUMonomer, an end-to-end deep-learning framework trained with input sequences spanning thousands of nucleotides on a joint RNA and single-stranded DNA dataset to predict nucleic acid structures directly from sequence. Despite requiring no auxiliary inputs, NUMonomer matches or outperforms leading prediction methods on benchmarks comprising CASP16 RNA targets and non-redundant sets of experimentally determined RNA and single-stranded DNA structures, with particularly pronounced improvements for longer RNAs. Its efficient and scalable architecture also reduces inference costs by approximately two orders of magnitude relative to the evaluated methods, enabling large-scale structure prediction. Together, these findings provide insight into generalization in biomolecular structure learning and establish NUMonomer as a practical framework for nucleic acid structure prediction.

## Introduction

Single-stranded nucleic acids perform diverse biological functions by adopting specific three-dimensional structures^1,2^. Despite decades of experimental effort, only a small fraction of known nucleic acid sequences have been characterized at atomic resolution^3–5^. Conserved sequence–structure relationships across diverse natural nucleic acids provide a foundation for computational structure prediction^6–8^. Accurate computational prediction could help bridge the widening gap between rapidly expanding sequence databases and limited structural data, while accelerating applications ranging from mechanistic studies to rational nucleic acid design^9,10^.

Conventional ab initio nucleic acid structure prediction approaches are guided by the thermodynamic principle that stable native conformations are associated with low free energy^11–14^. These methods explore conformational space using hand-crafted energy functions that combine physics-based terms with knowledge-based potentials derived from experimentally determined structures. These approaches can be effective for relatively simple folds but often struggle with structures stabilized by long-range, cooperative and context-specific interactions. Many methods, such as 3DRNA/DNA^15,16^, FARFAR2^17^ and SimRNA^11^, further restrict the conformational search space by incorporating fragment libraries, structural templates, base–base interactions inferred from multiple sequence alignments (MSAs), predicted secondary structures or other spatial restraints. However, the noise and uncertainty inherent in these auxiliary inputs can limit their effectiveness.

Deep learning has markedly advanced nucleic acid structure prediction. Compared with conventional approaches, deep learning models possess greater representational capacity and can automatically learn sequence–structure relationships from data, thereby potentially improving predictive generalization. The most prominent precedent comes from monomeric protein structure prediction, where deep learning methods have achieved accuracy approaching that of experimental structures for many proteins^18,19^. Inspired by this success, methods such as RoseTTAFoldNA^20^, DRFold^21^, DRFold2^22^, NUFold^23^, trRosettaRNA^24^, RhoFold+^25^, and AlphaFold3^26^ have extended deep learning frameworks to nucleic acid structure prediction by integrating sequence information with structure-related auxiliary inputs, including MSAs, secondary-structure annotations, and pretrained language model representations. These approaches have substantially improved predictive performance relative to many conventional methods, indicating that higher-capacity architectures combined with richer sequence–structure information can improve nucleic acid structure prediction. Nevertheless, accurate prediction remains challenging for numerous targets^27–29^, particularly long or structurally complex RNAs, despite the extensive use of available nucleic acid structures, structure-informed features, and diverse deep learning architectures in method development. Because the capacity of deep learning models can be readily increased by adding parameters, stacking layers, or incorporating additional modules, further progress is increasingly likely to depend on more reliable and diverse sequence–structure supervision rather than architectural complexity alone. From a computational perspective, the central challenge therefore lies in deriving more generalizable sequence–structure mappings from the limited set of experimentally determined nucleic acid structures currently available.

Experimentally determined nucleic acid structures remain underexploited across the folding hierarchy, from local conformations to global tertiary organization. At the global scale, nucleic acid architectures encode rich long-range and cooperative interactions that provide valuable supervision for deep learning models. However, input sequence-length constraints during training prevent current deep learning approaches from fully leveraging the global architecture of long nucleic acids. Eukaryotic 28S ribosomal RNA, which spans more than 4000 nucleotides^30^, exemplifies this limitation, as its global structure lies far beyond the sequence lengths used to train existing models. Consequently, global structural information remains only partially utilized, potentially limiting model generalization. Local structural information is likewise underutilized. Each experimentally determined structure comprises numerous local conformational patterns, many of which recur across diverse sequence and structural contexts^31,32^. This recurrence suggests that local sequence–structure relationships may be broadly generalizable. Although embedded within global folds, these local conformations represent distinct sequence–structure mappings that provide complementary supervision signals. Nevertheless, current deep learning models are trained on complete structures or fixed-length segments^20–22,24,25^, thereby limiting their ability to exploit reusable local conformational information. Together, experimentally determined structures encode rich multiscale information, spanning reusable local sequence–structure relationships to global folding organization. More effective utilization of these complementary supervision signals may substantially improve model generalization.

Cross-molecular relationships between RNA and single-stranded DNA (ssDNA) represent another potentially underexplored source of structural information. Although RNA and ssDNA differ in chemical composition and conformational preferences, both are single-stranded polynucleotides whose structures are governed by related physical interactions, including base pairing, base stacking, electrostatics, and backbone constraints. Prior studies have shown that sequence–structure relationships learned in one molecular context can transfer to related molecular systems. For example, deep learning models trained on monomeric proteins retain some ability to predict protein–protein complex structures^33–35^, suggesting that structural representations learned from monomeric proteins generalize to protein interfaces. Similarly, RNA secondary-structure prediction methods, such as MXFold2^36^, EternaFold^37^, and SPOT-RNA^38^, exhibit partial predictive capability for RNA–RNA interactions^39^, indicating that sequence–structure relationships learned from monomeric RNA may also transfer to intermolecular interfaces. However, current nucleic acid structure prediction methods are typically developed using RNA and ssDNA structural data separately. We therefore hypothesized that joint training on both molecular classes could improve prediction for each by exploiting shared structural representations. Although unified multimolecular frameworks such as AlphaFold3 demonstrate that multiple molecular classes can be represented within a common predictive framework, they do not establish whether RNA and ssDNA provide synergistic, independent, or potentially conflicting supervision during model training. Controlled joint-training experiments are therefore required to determine whether structural information from one nucleic acid class can improve prediction of the other.

We developed an end-to-end deep learning framework to investigate how multiscale structural information and cross-molecular relationships contribute to nucleic acid structure prediction. Its capacity to be trained with input sequences spanning thousands of nucleotides enables systematic investigation of the contribution of global fold topology in long nucleic acids. To assess intrinsic sequence–structure generalization while minimizing bias from external structural priors, the framework takes only the primary sequence as input and is trained exclusively on experimentally determined monomeric nucleic acid structures, without using model-predicted structures or auxiliary features such as MSAs and secondary-structure annotations. Within this framework, we trained a series of models under distinct training strategies to quantify the contributions of these sources of structural information. Benchmarks on recently released experimental RNA structures and RNA targets from the 16th Critical Assessment of Structure Prediction (CASP16)^40^ showed that dynamic length sampling over an expanded training sequence-length range substantially improves prediction accuracy, consistent with more effective utilization of structural information across multiple length scales. Furthermore, joint training on RNA and ssDNA structures substantially improved ssDNA structure prediction, supporting the existence of transferable sequence–structure relationships between the two molecular classes.

Guided by these findings, we developed NUMonomer, an end-to-end framework for single-sequence nucleic acid structure prediction. On recently released experimental RNA structures and CASP16 RNA targets, NUMonomer achieved a higher success rate than leading RNA structure prediction methods, including DRFold, DRFold2, trRosettaRNA, NUFold, RoseTTAFoldNA, RhoFold+, and AlphaFold3. To assess sequence-only performance, methods requiring MSAs were additionally evaluated with a single-row MSA containing only the query sequence. Under this setting, NUMonomer achieved a substantially higher success rate than all evaluated methods. For ssDNA, NUMonomer and AlphaFold3 achieved comparable success rates on recently released experimental structures, although AlphaFold3 may have benefited from exposure to a broader range of DNA-related structural information during training. Importantly, NUMonomer reduced inference cost by more than two orders of magnitude relative to the evaluated state-of-the-art methods and enabled structure prediction for RNAs of up to 5000 nucleotides with a peak GPU memory requirement of approximately 3 GB. Together, these results support multiscale and cross-molecular supervision as valuable strategies for improving generalization and establish NUMonomer as an accurate, efficient, and scalable framework for RNA and ssDNA structure prediction.

## Results

### Overview of NUMonomer

NUMonomer was trained exclusively on experimentally determined nucleic acid structures deposited in the Protein Data Bank (PDB)^5^. The overall data collection and filtering procedure is summarized in Fig. 1a. We extracted all nucleic acid chains from the PDB and applied quality filters, including a structural resolution better than 9 Å and a release date before 17 June 2023 for RNA. Because experimentally determined ssDNA structures are relatively scarce, we used an earlier release-date cutoff (17 June 2022) to reserve more recent ssDNA structures for model evaluation. Additional filtering criteria are described in the Methods. Because nucleic acid chains derived from macromolecular complexes may not retain stable intramolecular conformations in isolation, we further adopted the filtering strategy used in NUFold and removed chains lacking sufficient local contacts. The remaining RNA and ssDNA chains were clustered separately with CD-HIT^41^ at 80% sequence identity, yielding 847 RNA clusters and 299 ssDNA clusters. We randomly held out 20 RNA clusters and 20 ssDNA clusters as the validation set, whereas structures from the remaining clusters were used for training.

**Figure 1:**
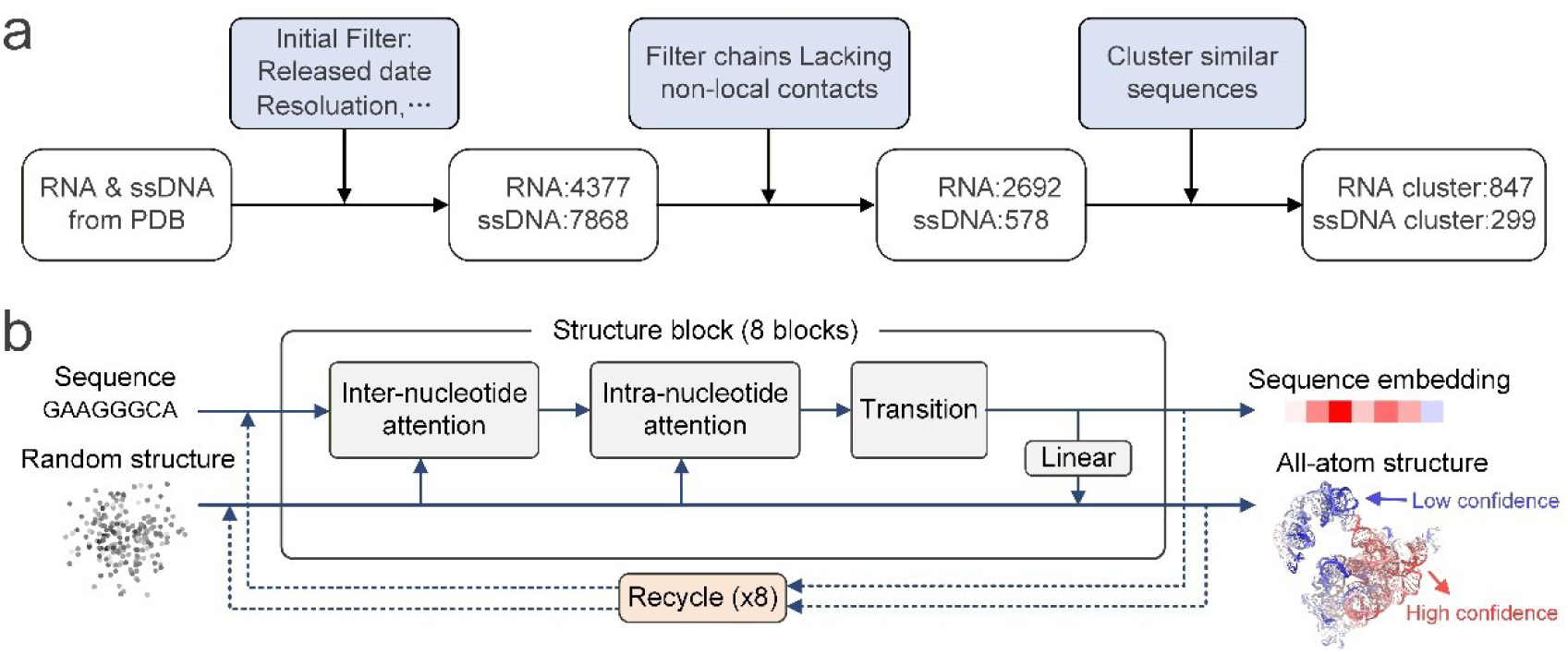
Overview of NUMonomer. **a,** Workflow for constructing the NUMonomer training dataset from experimentally determined RNA and ssDNA structures deposited in the Protein Data Bank (PDB). Blue boxes indicate data-processing operations. Numbers indicate the total number of sequences or clusters retained after each processing step. **b,** Architecture of NUMonomer for end-to-end nucleic acid structure prediction from primary sequence alone. Sequence representations and randomly initialized atomic coordinates are processed by a stack of eight structure blocks, each comprising Inter-nucleotide attention, Intra-nucleotide attention and Transition modules. The resulting sequence embeddings and all-atom structures are recycled eight times. Each structure block scales quadratically with sequence length. Details of the structure-block architecture are provided in the Supplementary Information.

NUMonomer employs an end-to-end deep learning strategy to predict three-dimensional all-atom nucleic acid structures and per-nucleotide confidence scores from a single sequence and randomly initialized atomic coordinates. As illustrated in Fig. 1b, the input sequence is first embedded into atom-level sequence representations, with each representation corresponding to a single atom within a nucleotide. Because each nucleotide contains multiple atom types, these representations are organized as a three-dimensional tensor spanning the inter-nucleotide, intra-nucleotide, and channel dimensions, thereby preserving atom-specific information across both sequence positions and nucleotide-internal atom types. NUMonomer performs structure prediction using a stack of eight structure blocks. Each structure block takes the current atom-level representations and atomic coordinates as input and outputs updated atom-level representations. The atom-level representations produced by the final structure block are then used to refine the atomic coordinates. To further improve prediction accuracy, we adopted the recycling strategy introduced in AlphaFold^18^, in which the predicted atomic coordinates and atom-level representations are recycled as inputs for subsequent refinement. After eight recycling iterations, the final atom-level representations and atomic coordinates are passed to a confidence module comprising four Evoformer layers, which predicts per-nucleotide Local Distance Difference Test^42^ (LDDT) scores. A detailed description of the NUMonomer workflow is provided in Supplementary Algorithm 1. Unlike triangle self-attention modules used in many nucleic acid structure prediction models, which scale cubically with sequence length, each NUMonomer structure block scales quadratically. Combined with the shallow architecture of only eight structure blocks, this computational efficiency enables training with input sequences of up to thousands of nucleotides and substantially reduces the computational cost of inference.

Consistent with the three-dimensional sequence representation, each structure block comprises an Inter-nucleotide attention module, an Intra-nucleotide attention module, and a transition module, which update atom-level representations along the sequence, intra-nucleotide, and channel dimensions, respectively. Inspired by the invariant point attention module in AlphaFold2, the inter-nucleotide attention module updates atom-level representations through self-attention conditioned on relative distances between corresponding atom types across different nucleotides, together with relative nucleotide positional information, thereby capturing inter-nucleotide structural interactions (the detailed architecture is shown in Fig. S1). The intra-nucleotide attention module adopts a similar architecture to update atom-level representations through self-attention conditioned on relative distances between atoms within each nucleotide, thereby modeling intra-nucleotide atomic interactions (the detailed architecture is shown in Fig. S2). Finally, the transition module updates atom-level representations through channel-wise feed-forward transformations, as illustrated in Fig. S3.

### Performance of NUMonomer in global RNA structure prediction

NUMonomer was first evaluated for RNA tertiary structure prediction on two non-redundant benchmark datasets: a set of recently determined experimental RNA structures, hereafter referred to as the Recent RNA dataset, and the publicly available RNA targets from CASP16, hereafter referred to as the CASP16 dataset. The Recent RNA dataset comprised 29 RNA structures deposited in the PDB between 17 June 2023 and 17 June 2024, with sequence lengths ranging from 6 to 4269 nucleotides. These structures were collected using the same procedure as that used to construct the NUMonomer training set but were filtered to ensure no greater than 80% sequence identity to any training RNA structure, as described in Methods. The CASP16 dataset comprised 16 RNA targets with sequence lengths ranging from 55 to 785 nucleotides, with corresponding experimental structures deposited in the PDB after June 2023, providing an independent benchmark for evaluating model generalization to previously unseen RNA folds. Structural agreement between predicted and experimental structures was assessed using TM-score^43^, which ranges from 0 to 1, with higher values indicating greater structural similarity.

Several state-of-the-art RNA structure prediction methods, including DRFold, DRFold2, NUFold, trRosettaRNA, RoseTTAFoldNA, RhoFold+, and AlphaFold3, were evaluated as baselines on the two benchmark datasets. These methods differ in their use of auxiliary inputs, including MSAs, predicted secondary structures, and pretrained language model representations, as well as in their architectures and training strategies, thereby providing a diverse set of reference methods. Except for DRFold2, all methods were evaluated on both benchmark datasets. DRFold2 was evaluated only on the CASP16 dataset because the Recent RNA dataset substantially overlaps with its training data. For methods requiring MSAs as input, alignments were generated by searching the NT^44^, RNAcentral^3^, and Rfam^4^ sequence databases using nhmmer^45^. Because RhoFold+, RoseTTAFoldNA, and DRFold support only limited input sequence lengths, longer targets were cropped to ensure compatibility. This preprocessing followed the sequence-cropping strategies used during the development of the respective methods.

Benchmarking NUMonomer and the reference methods using their standard inference settings showed that NUMonomer achieved global RNA structure prediction accuracy comparable to that of AlphaFold3 and higher than that of the other evaluated methods. On the Recent RNA dataset, NUMonomer achieved a mean TM-score of 0.431, compared with 0.441 for AlphaFold3; both methods outperformed the remaining methods by at least 0.14 (Fig. 2a), with detailed results provided in Table S1. A similar pattern was observed for the CASP16 dataset, for which NUMonomer and AlphaFold3 achieved mean TM-scores of 0.416 and 0.424, respectively, exceeding those of the other methods by approximately 0.10 (Fig. 2b), with detailed results provided in Table S1. DRFold2 was additionally evaluated on the CASP16 benchmark. Selecting the highest-scoring structure from 84 candidate conformations yielded a mean TM-score of 0.380, approaching that of NUMonomer. By contrast, the mean TM-score across individual DRFold2 candidates was 0.336, substantially lower than that of NUMonomer, as indicated by the shaded region in Fig. 2b. Notably, NUMonomer achieved comparable accuracy without relying on MSAs, predicted secondary structures, or other auxiliary features. To complement the mean TM-score analysis, we evaluated prediction success rates on the combined Recent RNA and CASP16 datasets, defining a successful prediction as one with a TM-score greater than 0.45. NUMonomer achieved a success rate of 40.0%, exceeding that of AlphaFold3 by 4.4 percentage points and those of all remaining methods by more than 20 percentage points, as shown in Fig. 2c. Together, these results indicate that NUMonomer generalizes to unseen RNA structures and that a sequence-only model can achieve global structural accuracy comparable to that of methods relying on auxiliary inputs.

**Figure 2:**
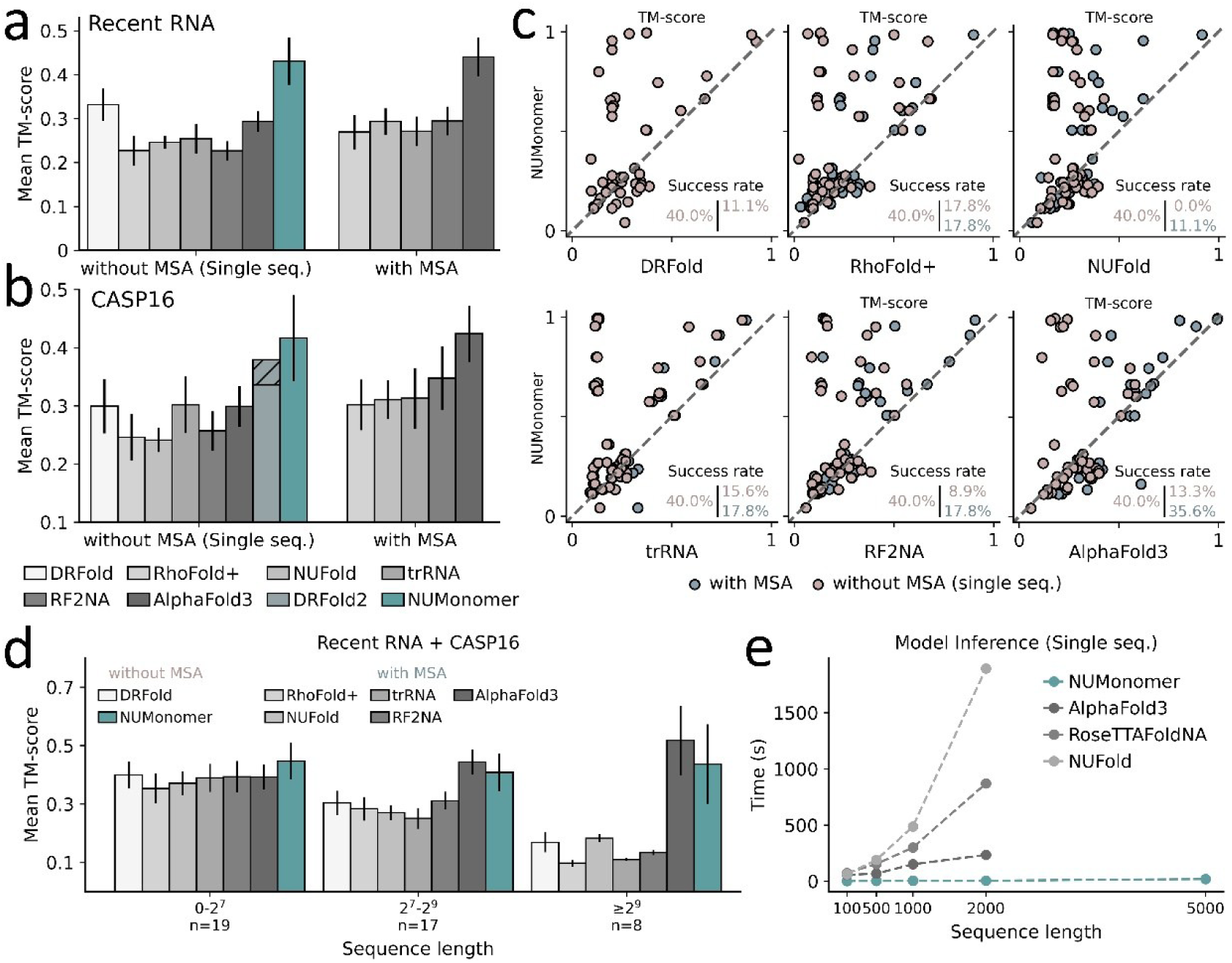
Performance of NUMonomer in global RNA structure prediction. **a-b,** Mean TM-scores of DRFold, NUMonomer, NUFold, RhoFold+, AlphaFold3, trRosettaRNA (trRNA), and RoseTTAFoldNA (RF2NA) on the Recent RNA **(a)** and CASP16 **(b)** datasets. DRFold2 was evaluated only on the CASP16 dataset; shaded regions indicate its performance after large-scale sampling. For MSA-independent methods, including DRFold, DRFold2, and NUMonomer, results are shown in the without MSA group; for MSA-dependent methods, including NUFold, RhoFold+, AlphaFold3, trRosettaRNA, and RoseTTAFoldNA, results obtained under their standard inference settings are shown in the with MSA group, whereas results obtained under single-sequence inference settings are shown in the without MSA group. The Recent RNA and CASP16 datasets contain 29 and 16 RNA structures, respectively. Black vertical lines indicate the standard deviation across RNA targets. **c,** Head-to-head comparison of TM-scores between NUMonomer and each benchmark method on the combined Recent RNA and CASP16 datasets. Percentages denote the success rate at a TM-score threshold of >0.45. For MSA-dependent methods, pinkish grey and bluey grey denote results obtained under single-sequence and standard inference settings, respectively. **d,** Mean TM-scores of NUMonomer, NUFold, RhoFold+, AlphaFold3, trRosettaRNA, DRFold, and RoseTTAFoldNA benchmarked under their standard inference settings across RNA sequence-length groups in the combined Recent RNA and CASP16 datasets. Black vertical lines indicate the standard deviation across RNA targets. n values indicate the number of RNA structures in each interval. **e,** Inference times of NUFold, RoseTTAFoldNA, AlphaFold3 and NUMonomer for RNA sequences of different lengths (measured in nucleotides) under the single-sequence setting on a single NVIDIA H100-PCIe GPU.

Given that accurate sequence-to-structure mapping is important for RNA structure design and other high-throughput downstream applications, we next evaluated the sequence-only prediction performance of each method on the Recent RNA and CASP16 datasets. For methods requiring MSAs, the alignments were reduced to a single row containing only the query sequence. The resulting mean TM-scores for each method are summarized in Table S1. Except for trRosettaRNA, removal of homologous-sequence information reduced the accuracy of all MSA-dependent methods, thereby widening the performance gap between NUMonomer and the reference methods in the sequence-only setting (Fig. 2b). The limited effect on trRosettaRNA is consistent with the original study, which reported only weak dependence on MSA depth^24^. Prediction success rates showed a similar trend, declining for MSA-dependent methods when only the query sequence was provided (Fig. 2c). Most MSA-dependent methods showed reductions of approximately 0.05 in mean TM-score, whereas AlphaFold3 showed substantially larger decreases of 0.17 on the Recent RNA dataset and 0.13 on the CASP16 dataset. Length-stratified analysis indicated that the decrease in AlphaFold3 performance was largely confined to longer RNA targets, for which its accuracy declined substantially in the absence of homologous-sequence information (Fig. S4; compare Fig. 2d with Fig. S5). By contrast, the other reference methods already showed relatively low accuracy for longer RNAs under their standard MSA-enabled settings and therefore underwent only small additional decreases after removal of homologous sequences (Fig. 2d and Fig. S5).

Beyond the aggregate benchmark statistics, length-stratified analysis further showed that NUMonomer maintained competitive accuracy across diverse RNA sequence lengths, indicating robust generalization across target sizes. For this analysis, structures from the Recent RNA and CASP16 datasets were pooled and stratified into three length bins: short RNAs of no more than 128 nucleotides, medium-length RNAs of 129–512 nucleotides and long RNAs of more than 512 nucleotides. Performance within each group was assessed using the mean TM-score of predictions generated under the standard inference setting for each method. As shown in Fig. 2d (detailed results provided in Table S2), NUMonomer achieved the highest mean TM-score on short RNAs, outperforming all reference methods by approximately 0.05, and maintained a pronounced advantage over most methods on longer RNAs. AlphaFold3 was the only exception, achieving slightly higher mean TM-scores than NUMonomer for the medium- and long-length targets. Notably, all methods except NUMonomer and AlphaFold3 exhibited a pronounced decline in accuracy as RNA length increased. Given that NUMonomer and AlphaFold3 were trained on longer nucleic acid sequences than the other evaluated methods, this pattern is consistent with the possibility that longer RNA chains contain structural features that are underrepresented in shorter sequences and that exposure to longer sequence contexts during training may facilitate learning of these features.

Inference-time benchmarking further demonstrated the computational efficiency and scalability of NUMonomer. To standardize the comparison, all methods were evaluated using single-sequence inputs on a single NVIDIA H100 PCIe GPU. The reported runtimes included only model inference and excluded molecular-dynamics relaxation and input preparation steps, such as secondary-structure prediction and MSA searches. Consequently, these measurements may underestimate the end-to-end runtimes of methods that require substantial preprocessing^45,46^. DRFold, DRFold2, and trRosettaRNA were excluded because their inference pipelines rely on CPU-based structure assembly, which can substantially increase runtime. Previous evaluations have shown that these methods are considerably slower than AlphaFold3^22,28,29^. RhoFold+ was also excluded because its Python environment was incompatible with the H100 configuration used in this analysis. Among the methods evaluated under the standardized setting, NUMonomer was approximately two orders of magnitude faster across the tested sequence lengths (Fig. 2e), with detailed results provided in Table S3. Moreover, by adjusting inference parallelism, NUMonomer processed RNAs of up to 5,000 nucleotides with a peak GPU memory requirement of approximately 3 GB, whereas the other benchmarked methods encountered out-of-memory errors for sequences of this length. Together with its predictive accuracy, these results establish NUMonomer as a computationally efficient and scalable framework for sequence-based RNA structure prediction.

### Local RNA geometry and nucleotide interaction networks predicted by NUMonomer

To assess the local structural fidelity of NUMonomer and the reference methods, predictions for targets in the combined Recent RNA and CASP16 datasets were evaluated using the mean of circular quantities (MCQ)^47^ and interaction network fidelity (INF)^48,49^. MCQ quantifies differences in torsion-angle geometry between predicted and experimental structures, with lower values indicating closer structural agreement. Specifically, MCQ measures deviations in the α, β, γ, δ, ε, ζ, and χ torsion angles^50^. INF quantifies the recovery of nucleotide interaction networks in the predicted structures and was calculated using the RNA_assessment^48^ package. The analysis included Watson–Crick base pairs, non-Watson–Crick base pairs, and base-stacking interactions. INF ranges from 0 to 1, with higher values indicating greater agreement with the corresponding experimental structure.

NUMonomer accurately predicted local RNA torsion-angle geometry. Figure 3a shows the distributions of MCQ values for structures predicted by NUMonomer and the reference methods on the combined Recent RNA and CASP16 datasets using their standard inference settings. MSA-dependent methods were additionally evaluated using single-sequence alignments containing only the query sequence. Except for RhoFold+ and trRosettaRNA, the evaluated methods produced broadly similar MCQ distributions, even when the MSA-dependent methods were evaluated without homologous-sequence information. The substantial overlap among these distributions may reflect the tendency of RNA torsion angles to cluster into a limited number of dominant conformational states^51^, enabling models that recover these recurring states to achieve relatively low MCQ values. Examples of failed predictions from each method are shown in Fig. S6. Nevertheless, NUMonomer achieved the lowest median MCQ of 17.1 among the methods evaluated using single-sequence or standard inference settings, with detailed results provided in Table S4. To directly compare NUMonomer and DRFold2, we conducted a head-to-head analysis of their predictions for the CASP16 RNA targets, with DRFold2 evaluated using its standard 84-candidate sampling protocol. NUMonomer achieved lower MCQ values than DRFold2 for all targets (Fig. 3b), indicating greater local torsion-angle accuracy.

**Figure 3:**
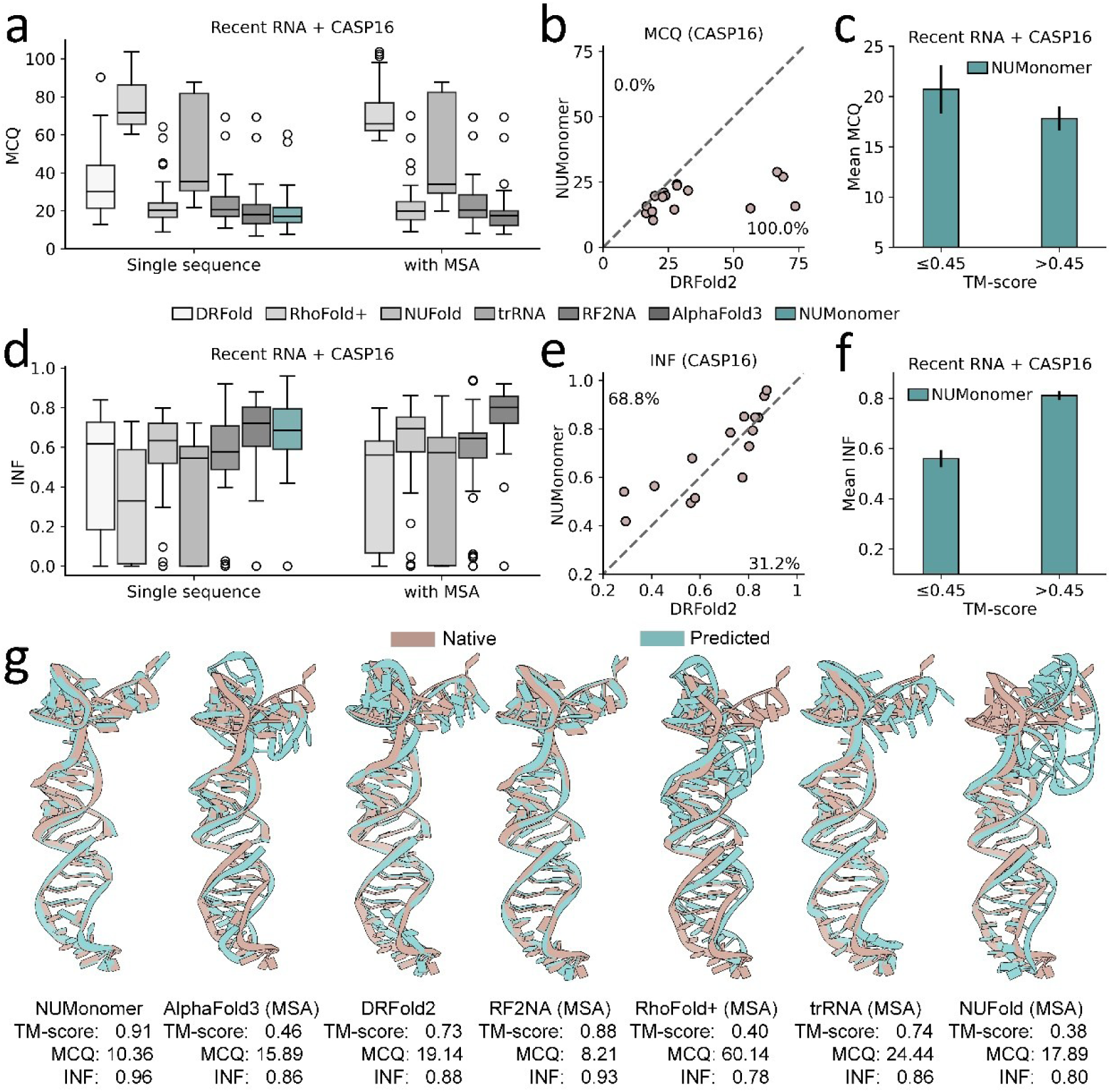
Local quality of RNA structures predicted by NUMonomer. **a,** Distribution of MCQ scores for structures predicted by NUMonomer, NUFold, RhoFold+, DRFold, trRosettaRNA (trRNA), RoseTTAFoldNA (RF2NA), and AlphaFold3 on the combined Recent RNA and CASP16 datasets. For MSA-independent methods (DRFold and NUMonomer), results are shown in the Single sequence group; for MSA-dependent methods (NUFold, RhoFold+, AlphaFold3, trRosettaRNA, and RoseTTAFoldNA), results obtained under their standard inference settings are shown in the With MSA group, whereas results obtained under single-sequence inference settings are shown in the Single sequence group. Box plots show the 25–75% interquartile range (box limits) and the median (center line). **b,** Head-to-head comparison of MCQ scores between NUMonomer and DRFold2 predictions on the CASP16 dataset. Percentages indicate the proportion of targets in the upper and lower triangular regions. **c,** Mean MCQ scores of NUMonomer predictions grouped by whether the TM-score is >0.45 on the combined Recent RNA and CASP16 datasets. Black vertical lines indicate the standard deviation across RNA targets. **d,** Distribution of INF scores for structures predicted by NUMonomer, NUFold, RhoFold+, DRFold, trRosettaRNA (trRNA), RoseTTAFoldNA (RF2NA), and AlphaFold3 on the combined Recent RNA and CASP16 datasets. For MSA-independent methods (DRFold, and NUMonomer), results are shown in the Single sequence group. For MSA-dependent methods (NUFold, RhoFold+, AlphaFold3, trRosettaRNA, and RoseTTAFoldNA), results obtained under their standard inference settings are shown in the with MSA group, whereas results obtained under single-sequence inference settings are shown in the single sequence group. **e,** Head-to-head comparison of INF scores between NUMonomer and DRFold2 predictions on the CASP16 dataset. Percentages indicate the proportion of targets in the upper and lower triangular regions. **f,** Mean INF scores of NUMonomer predictions grouped by whether the TM-score is >0.45 on the combined Recent RNA and CASP16 datasets. Black vertical lines indicate the standard deviation across RNA targets. **g,** Structures predicted by NUMonomer and the reference methods for CASP16 target R1203 using their standard inference settings.

NUMonomer also recovered local nucleotide-interaction networks with high accuracy. Fig. 3d shows the distributions of INF values for predictions generated by NUMonomer and reference methods on the combined Recent RNA and CASP16 datasets under their standard inference settings. Methods requiring MSAs were additionally evaluated using single-row alignments containing only the query sequence. Under the standard inference settings, NUMonomer achieved a slightly lower median INF value than AlphaFold3 and NUFold but higher median INF values than the other reference methods (detailed results provided in Table S5). Under the single-sequence setting, the MSA-dependent methods exhibited modest decreases in INF (detailed results provided in Table S5). We also conducted a head-to-head INF comparison of NUMonomer and DRFold2 on CASP16 targets predicted by both methods, with DRFold2 evaluated using its standard 84-candidate sampling protocol. As shown in Fig. 3e, NUMonomer achieved higher INF values than DRFold2 for 68.8% of all targets, indicating greater overall accuracy in recovering nucleotide interactions. Specifically, NUMonomer achieved a mean INF of 0.697, compared with 0.636 for DRFold2.

NUMonomer predictions with higher global structural accuracy tended to exhibit greater local structural fidelity. Predictions for targets in the Recent RNA and CASP16 datasets were divided into two groups: 27 failed predictions with TM-scores of 0.45 or lower and 18 successful predictions with TM-scores greater than 0.45. Successful predictions had a lower mean MCQ than failed predictions (17.8 versus 20.7; Fig. 3c). The association between INF and TM-score was examined in the same manner, as shown in Fig. 3f. Successful predictions had a higher mean INF than failed predictions, with values of 0.81 and 0.56, respectively. Together, these analyses indicate that greater global fold accuracy in NUMonomer predictions is associated with improved local torsion-angle geometry and nucleotide-interaction fidelity. We further illustrated the concordance among the three metrics using a representative CASP16 target (Fig. 3g). For this target, NUMonomer achieved a higher TM-score than the comparison method, together with more favorable local-accuracy metrics, namely a lower MCQ and a higher INF.

### Performance of NUMonomer in ssDNA structure prediction

For ssDNA tertiary structure prediction, NUMonomer was evaluated on a benchmark comprising 21 recently determined ssDNA structures, hereafter referred to as the Recent ssDNA dataset. The structures ranged from 16 to 53 nucleotides in length and were deposited in the PDB between 17 June 2022 and 17 June 2024. We curated the dataset using the protocol applied to construct the NUMonomer training set and excluded targets sharing more than 80% sequence identity with any ssDNA sequence in the training set, as described in Methods. Because the ssDNA targets were relatively short (Fig. S7), global agreement between the predicted and experimental structures was assessed using root-mean-square deviation (RMSD) calculated over C3′ atoms rather than TM-score, with lower RMSD values indicating closer structural agreement. Local structural fidelity was evaluated using MCQ and INF, which were calculated following the same procedures used for the predicted RNA structures. Given the scarcity of dedicated ssDNA structure prediction methods and the competitive performance of AlphaFold3 in previous benchmarks^52–54^, AlphaFold3 was included as the sole reference method.

NUMonomer and AlphaFold3 showed comparable global ssDNA structure prediction performance. The distributions of RMSD values obtained by the two methods on the Recent ssDNA dataset are shown in Fig. 4a, with detailed results provided in Table S6. NUMonomer achieved a median RMSD of 8.20 Å, compared with 7.47 Å for AlphaFold3. Consistent with this result, when successful predictions were defined by a relaxed RMSD threshold of 6 Å, the two methods achieved similar success rates of 38% for NUMonomer and 42.8% for AlphaFold3. These relatively low success rates underscore the continuing difficulty of reliable ssDNA tertiary structure prediction. Target-level analysis further showed that, for many structures in the Recent ssDNA dataset, NUMonomer yielded RMSD values similar to those obtained with AlphaFold3 (Fig. 4b). However, AlphaFold3 achieved substantially lower RMSD values for three targets. Sequence-similarity analysis indicated that these targets had close matches among experimentally determined double-stranded DNA (dsDNA) structures (Fig. 4c). Given that dsDNA structures were included in the AlphaFold3 training data but not in that of NUMonomer, these results suggest that representations learned from dsDNA may be partially transferable to ssDNA and may have contributed to AlphaFold3’s improved performance on these targets. Overall, NUMonomer achieved a lower RMSD than AlphaFold3 for 11 of 21 targets (52.4%), whereas AlphaFold3 achieved a lower RMSD for the remaining 10 targets (47.6%).

**Figure 4:**
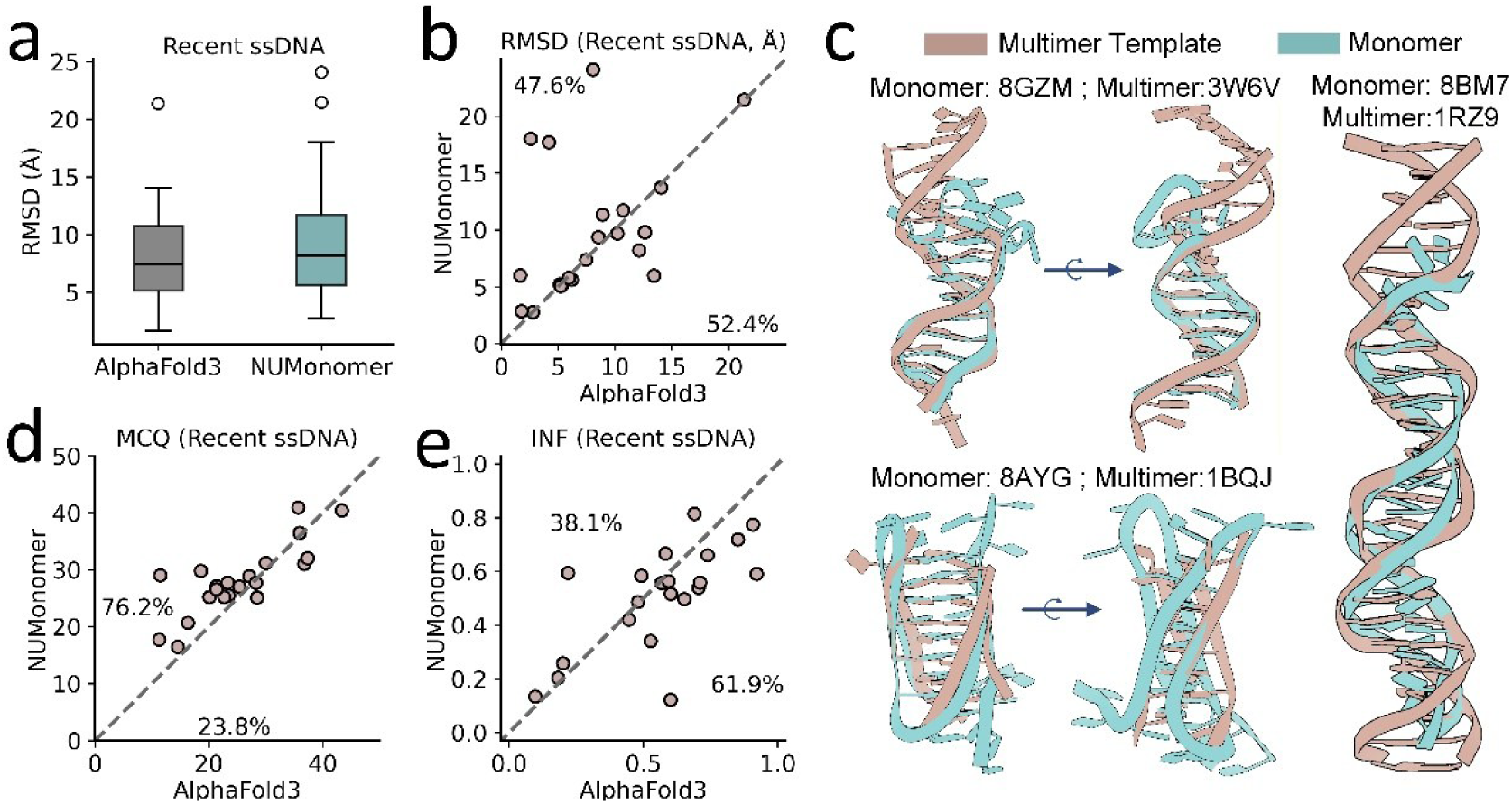
Performance of NUMonomer in ssDNA structure prediction. **a,** Distribution of RMSD values for structures predicted by NUMonomer and AlphaFold3 on the Recent ssDNA dataset. The Recent ssDNA dataset contains 21 ssDNA structures. Box plots show the interquartile range (25th–75th percentiles; box limits) and the median (center line). **b,** Head-to-head comparison of RMSD values between NUMonomer and AlphaFold3 predictions on the Recent ssDNA dataset. Percentages indicate the proportion of targets in the upper and lower triangular regions. **c,** Representative examples showing structural similarity between selected ssDNA targets and experimentally determined dsDNA structures. The labels following each colon indicate the corresponding PDB IDs. **d-e,** Head-to-head comparison of MCQ (**d**) and INF (**e**) values between NUMonomer and AlphaFold3 predictions on the Recent ssDNA dataset. Percentages indicate the proportion of targets in the upper and lower triangular regions.

Local geometry and nucleotide-interaction networks in structures predicted by NUMonomer and AlphaFold3 on the Recent ssDNA dataset were also evaluated using MCQ and INF. Pairwise comparisons of MCQ and INF values between the two methods are shown in Fig. 4d,e, with detailed results provided in Table S6. Overall, NUMonomer exhibited slightly lower local structural fidelity than AlphaFold3. Specifically, NUMonomer achieved a median MCQ of 27.7 and a median INF of 0.557, while AlphaFold3 achieved a median MCQ of 23.5 and a median INF of 0.593. Together with the global structural results, these findings indicate that reliable ssDNA structure prediction remains challenging at both global and local scales. Given the substantial differences between NUMonomer and AlphaFold3 in model architecture, training strategies, and training datasets, the challenges observed for both methods may partly reflect broader limitations in ssDNA structure prediction, including the conformational flexibility of ssDNA and the limited number and diversity of experimentally determined ssDNA structures.

### Ablation studies

We developed a series of ablation models to quantify the contributions of key training and inference strategies in NUMonomer, including clash and bond losses, dynamic length sampling, fine-tuning on an extended sequence-length range, inference-time sampling, and joint RNA-ssDNA training. Unless otherwise specified, all ablation models were trained on the same dataset as NUMonomer and implemented using the same deep learning framework. Performance was evaluated on the combined CASP16 and Recent RNA datasets, with TM-score used to assess global RNA structure prediction, and on the Recent ssDNA dataset, with RMSD used to assess global ssDNA structure prediction.

Clash and bond losses improved nucleic acid structure prediction. Such geometric losses have been widely used during training or post-training refinement of protein and RNA structure prediction models to enhance the physical plausibility of generated structures and can also modestly improve global structural accuracy^25,55,56^. However, their utility for ssDNA structure prediction remains unclear. We first constructed a baseline model, denoted as Baseline, which followed a conventional training configuration with a fixed crop length of 384 nucleotides and did not use dynamic length sampling, fine-tuning on an extended length range, or inference-time sampling. To assess the contribution of geometric losses, we trained an additional ablation model based on this Baseline model but without the clash and bond losses, referred to as the “-clash & bond losses” model. The performance of the two models on the RNA and ssDNA benchmark datasets is shown in Fig. 5a, with detailed results provided in Table S7. Removing the clash and bond losses reduced prediction accuracy, decreasing the median TM-score for RNA by 0.05 and reducing overall ssDNA structure prediction performance (see Fig. S8). These results indicate that clash and bond losses improve structure prediction for both RNA and ssDNA, supporting the broader applicability of these geometric constraints in biomolecular structure prediction.

**Figure 5:**
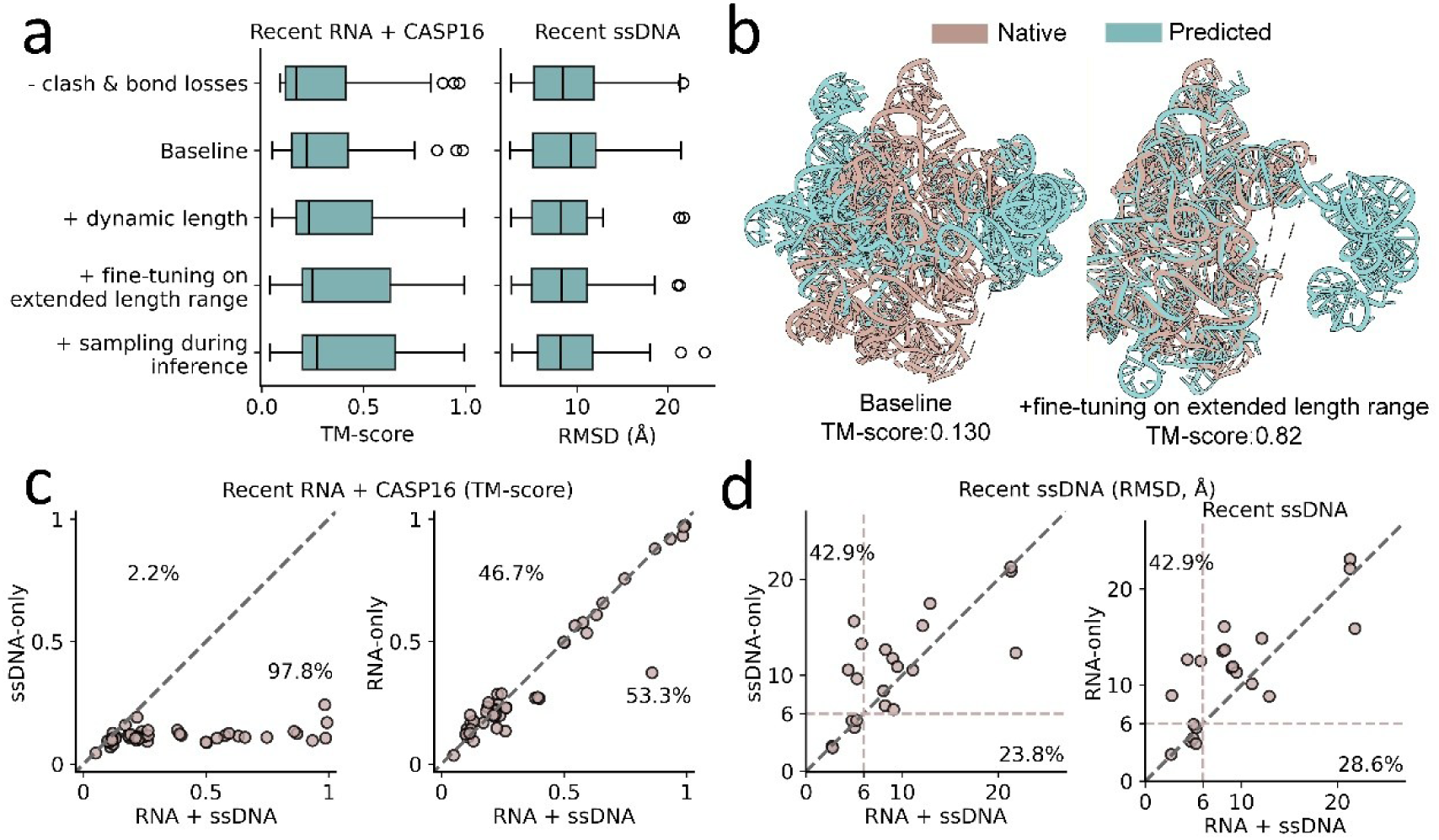
Ablation analysis and interpretation of NUMonomer. **a,** Performance of the ablation models on the combined Recent RNA and CASP16 datasets and on the Recent ssDNA dataset. RNA structures were evaluated using TM-score, whereas ssDNA structures were evaluated using RMSD. Box plots show the interquartile range (25th–75th percentiles; box limits) and the median (center line). **b,** Predictions generated by the baseline model (TM-score, 0.130) and the model fine-tuned on an extended sequence-length range (TM-score, 0.820) for a 621-nucleotide RNA target (PDB 8K15). **c,** Head-to-head comparison of models trained on RNA structures only, ssDNA structures only, or a mixture of RNA and ssDNA structures on the combined Recent RNA and CASP16 datasets. Performance was evaluated using TM-score. Percentages indicate the proportion of targets in the upper and lower triangular regions. **d,** Pairwise comparison of models trained exclusively on RNA structures, exclusively on ssDNA structures, or jointly on RNA and ssDNA structures, evaluated on the Recent ssDNA dataset. Performance was assessed using RMSD. Percentages indicate the proportion of targets with RMSD values below 6 Å.

Learning hierarchical structural organization from experimentally determined structures through dynamic length sampling over an extended length range improved global RNA structure prediction. To evaluate the contribution of this strategy, we introduced a dynamic length sampling strategy into the Baseline model by randomly cropping each experimentally determined structure to a length sampled between 16 and 2048 nucleotides. The resulting model was denoted the “+dynamic length” model. To further assess whether training on longer sequence lengths improves generalization, we fine-tuned the “+dynamic length” model using an expanded dynamic length range from 16 to 2560 nucleotides, denoted as “+ fine-tuning on extended length range” model. Performance comparisons on the RNA and ssDNA benchmark datasets showed that dynamic length sampling over the extended length range improved global RNA structure prediction, but had only a limited effect on ssDNA (Fig. 5a and Fig. S9; detailed results provided in Table S7). This difference is likely explained by the sequence-length distributions of the training data. Whereas a substantial fraction of RNA structures in the NUMonomer training dataset are relatively long, most ssDNA structures are considerably shorter (Fig. S10). Consequently, extending the dynamic length range exposes the model to substantially more diverse sequence–structure relationships for RNA, but provides comparatively little additional information for ssDNA. The benefit of training on longer sequence lengths is further illustrated in Fig. 5b. For a 621-nucleotide RNA target, the Baseline model achieved a TM-score of only 0.13, whereas the “+ fine-tuning on extended length range” model achieved a TM-score of 0.82.

Given that the designed deep learning framework generates structures from randomly initialized coordinates and predicts a confidence score for each output, we next investigated whether inference-time sampling could further improve predictive performance. In this ablation, the “+ fine-tuning on extended length range” model generated ten candidate structures for each sequence, and the candidate with the highest predicted LDDT (pLDDT) score was retained as the final prediction. This strategy modestly improved both RNA and ssDNA structure prediction. As shown in Fig. 5a, the “+ fine-tuning on extended length range” model achieved a median TM-score of 0.247 for RNA and a median RMSD of 8.29 Å for ssDNA; inference-time sampling increased the median RNA TM-score to 0.270 and reduced the median ssDNA RMSD to 8.20 Å. We next investigated whether the modest gains were limited by insufficient structural diversity among the sampled candidates or by imperfect confidence-based ranking. To estimate the upper bound of the improvement attainable from the sampled candidate pool, we applied an oracle selection strategy that retained the candidate with the highest TM-score for RNA or the lowest RMSD for ssDNA. Compared with selection based on pLDDT, oracle selection further increased the median RNA TM-score by 0.038 and reduced the median ssDNA RMSD by 0.5 Å (Fig. S11). These results indicate that inference-time sampling enables NUMonomer to generate higher-quality structural candidates, but that the current confidence metric does not consistently identify the best-performing candidate. Improved confidence estimation or candidate-ranking strategies could therefore further increase the benefits of inference-time sampling.

Finally, we assessed whether joint training on RNA and ssDNA structures improved performance relative to training on either molecular class alone. Starting from the Baseline configuration, we trained two additional ablation models exclusively on RNA or ssDNA structures, designated RNA-only and ssDNA-only, respectively. For this analysis, the original Baseline model, trained on both RNA and ssDNA structures, was designated RNA + ssDNA. The ssDNA-only model generalized poorly to RNA targets, whereas the RNA-only and RNA + ssDNA models achieved comparable performance (Fig. 5c), indicating that including ssDNA structures did not materially affect RNA structure prediction. In contrast, the RNA-only model retained measurable predictive capability on ssDNA targets, and the RNA + ssDNA model consistently outperformed the ssDNA-only model (Fig. 5d). Together, these findings indicate asymmetric transfer between the two molecular classes. Given the substantially larger number of experimentally determined RNA structures available for training (Fig. S10), this asymmetry suggests that RNA and ssDNA share transferable structural patterns, whereas the limited availability of ssDNA structures constrains reciprocal transfer to RNA prediction.

## Discussion

Accurate nucleic acid structure prediction remains a longstanding challenge in computational biology. Within the deep learning paradigm, the incorporation of structure-informed auxiliary features has driven much recent progress in this field. Our results suggest a complementary route in which experimentally determined structures themselves encode rich and reusable sequence–structure relationships across multiple hierarchical scales, from local conformational patterns to global topologies. Dynamic sampling across an expanded range of sequence lengths during model training enables more effective use of these intrinsic multiscale supervision signals, thereby improving the generalization and predictive performance of nucleic acid structure prediction models.

Our results further indicate that RNA and ssDNA share transferable structural information that can improve nucleic acid structure prediction. Although these molecules differ in chemical composition and conformational preferences, both are single-stranded polynucleotides shaped by related physical constraints, including base pairing, base stacking, electrostatics and backbone geometry. The improved performance achieved by joint RNA–ssDNA training relative to training on either molecular class alone suggests that these shared structural regularities provide useful cross-molecular supervision. Notably, this transfer was asymmetric: RNA structures improved ssDNA structure prediction, whereas ssDNA structures provided little detectable benefit for RNA structure prediction, likely reflecting the substantially smaller number and shorter lengths of experimentally determined ssDNA structures available for training.

Guided by these findings, we developed NUMonomer, an end-to-end framework for nucleic acid structure prediction from primary sequence alone. In addition to learning hierarchical structural organization and shared RNA–ssDNA representations, NUMonomer incorporates geometric loss terms that penalize steric clashes and bond-geometry violations, together with inference-time sampling to further improve structural plausibility and prediction robustness. Clash and bond losses have previously been used in protein and RNA structure prediction to improve the physical realism of predicted structures and have been shown to modestly improve global structural accuracy. Our results show that these geometric constraints also benefit ssDNA prediction. Inference-time sampling, in which multiple candidate structures are generated and ranked using the predicted confidence score, provided additional but modest gains. However, the substantially better performance achieved by oracle selection suggests that more accurate confidence estimation and candidate ranking represent promising directions for future development. Across benchmarks comprising CASP16 RNA targets and recently released experimental RNA structures, NUMonomer substantially outperformed leading RNA structure prediction methods, including DRFold, DRFold2, trRosettaRNA, NUFold, RoseTTAFoldNA and RhoFold+. NUMonomer achieved performance comparable to AlphaFold3 when AlphaFold3 was provided with MSAs, while substantially outperforming it under the single-sequence setting. On recently released experimental ssDNA structures, NUMonomer achieved a success rate comparable to AlphaFold3, although the limited absolute accuracy of both methods indicates that accurate ssDNA structure prediction remains challenging.

Future improvements to NUMonomer may be achieved by introducing auxiliary features and incorporating experimentally determined nucleic acid complexes as training data. Although the current model achieves competitive performance without auxiliary inputs, the optional integration of reliable multiple sequence alignments, secondary-structure annotations and pretrained language-model representations may further improve accuracy for challenging targets, particularly RNAs with novel folds or extensive long-range interactions. Building on the transferable structural representations shared between RNA and ssDNA, future models may also benefit from training on a broader range of experimentally determined nucleic acid complexes, including RNA–RNA interactions, DNA–DNA assemblies and RNA–DNA complexes. Such structural data may provide transferable supervision for monomeric nucleic acid prediction, and integrating these structures during model training may further improve both prediction accuracy and model generalization.

## Methods

### Training and validation datasets

We retrieved all nucleic acid chains deposited in the Protein Data Bank (PDB) prior to 17 June 2023 to construct the NUMonomer training dataset. Chains were retained according to the following criteria: (1) a reported resolution better than 9 Å, when available; (2) release before 17 June 2023 for RNA chains and before 17 June 2022 for ssDNA chains, with the earlier ssDNA cutoff used to reserve more recent structures for model evaluation; (3) inclusion of only standard nucleotides; (4) at least four resolved residues; and (5) at least four non-local contacts. A non-local contact was defined as a minimum heavy-atom distance of <5 Å between nucleotides *i* and *j*, where |*j*-*i*| > 3. Chains with identical sequences were grouped, yielding 2692 unique RNA sequences and 578 unique ssDNA sequences. For sequences associated with multiple experimental structures, the structure with the greatest number of non-local contacts was selected for model training. The remaining RNA and ssDNA chains were clustered separately with CD-HIT^41^ at 80% sequence identity (-c 0.8 -M 0 -T 0 -g 1 -n 2 -d 60 -G 1 -r 0 -l 1), resulting in 847 RNA clusters and 299 ssDNA clusters. We randomly selected 20 clusters from each molecular class for validation, and used structures from the remaining clusters for training.

### Recent RNA dataset

RNA chains deposited in the PDB between 17 June 2023 and 17 June 2024 were used to construct the Recent RNA dataset. After applying the same filtering criteria as for the training and validation datasets and grouping structures with identical sequences, 129 unique RNA sequences were obtained. For sequences associated with multiple experimental structures, the structure containing the greatest number of non-local contacts was selected for benchmark construction. The remaining 129 chains were clustered against the combined training and validation dataset using CD-HIT-EST-2D (-i2 test.fasta -i train+valid.fasta -c 0.8 -M 0 -T 0 -g 1 -n 2 -d 60 -G 1 -r 0 -l 1) at 80% sequence identity to remove redundant sequences. This procedure yielded a benchmark dataset comprising 29 unique RNA chains.

### CASP16 dataset

All 16 publicly available RNA targets from https://predictioncenter.org/casp16/targetlist.cgi were collected to construct the CASP16 dataset. The CASP16 targets were released after the construction of the NUMonomer training dataset, thereby effectively eliminating the possibility of data leakage. The dataset comprised the following targets: R1203, R1205, R1209, R1211, R1212, R1242, R1251, R1260, R1261, R1262, R1263, R1264, R1283v1, R1285, R1286, and R1296.

### Recent ssDNA dataset

ssDNA chains deposited in the PDB between 17 June 2022 and 17 June 2024 were used to construct the Recent ssDNA dataset. After applying the same filtering criteria as for the training and validation datasets and grouping structures with identical sequences, 51 unique ssDNA sequences were obtained. For sequences associated with multiple experimental structures, the structure containing the greatest number of non-local contacts was selected for benchmark construction. The remaining 51 chains were clustered against the combined training and validation dataset using CD-HIT-EST-2D (-i2 test.fasta -i train+valid.fasta -c 0.8 -M 0 -T 0 -g 1 -n 2 -d 60 -G 1 -r 0 -l 1) at 80% sequence identity to remove redundant sequences. This procedure yielded a benchmark dataset comprising 21 unique ssDNA chains.

### Training protocol

During training, nucleic acid structures were randomly sampled from the training dataset according to weights determined by nucleic acid type, sequence length, and cluster size. RNA structures and RNA sequence clusters substantially outnumbered their ssDNA counterparts, and RNA sequences were generally longer than ssDNA sequences. To reduce the overrepresentation of large clusters while assigning greater sampling weights to longer sequences, the unnormalized sampling weights for each RNA and ssDNA structure were defined as 2×min(max(L, 32), 512)/C and min(max(L, 32), 512)/C, respectively, where L is the sequence length and C denotes the number of sequences in the cluster to which the corresponding nucleic acid structure belongs.

For each sampled nucleic acid structure, an upper bound on sequence length, denoted by *L_upper_*, was uniformly sampled from the integers between 16 and 2048 during initial training and between 16 and 2560 during fine-tuning. The input sequence length was defined as min(*L_upper_*, L), where L denotes the original sequence length. During training, positions in each input sequence were randomly selected for masking, with the masking rate uniformly sampled between 0% and 20%. Of the selected positions, 80% were replaced with a special mask token, 10% with randomly selected nucleotides, and 10% were left unchanged. During fine-tuning, the masking-rate range was expanded to 0–50%.

NUMonomer and all ablation models were trained on NVIDIA H100 GPUs using automatic mixed precision, an exponential moving average of model parameters, gradient accumulation, and activation checkpointing for each recycle iteration and structure block. Model parameters were optimized using AdamW^57^ with a learning rate of 0.001 and a weight decay of 0.001. Training was terminated when the mean LDDT on the validation set converged. For the RNA-only and ssDNA-only models, the corresponding validation sets contained only RNA and ssDNA structures, respectively. The batch size was 24 for NUMonomer and 4 for each ablation model. During training, each input sequence was processed through eight recycling iterations, yielding eight candidate structures. To expose the confidence module to a more diverse set of predicted structures, one of the eight candidates was randomly selected for confidence-module training. To improve training stability, per-sample gradients were clipped to a global norm of 0.1.

The training objective of NUMonomer comprised five terms: clash loss, bond loss, pLDDT loss, weighted frame-aligned point error (FAPE) loss, and distance root-mean-square deviation (dRMSD) loss. The clash loss penalizes steric clashes by penalizing predicted interatomic distances below a reference-dependent lower bound: 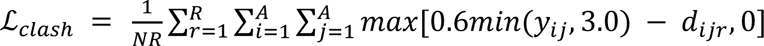, where N is the number of nucleotides, A is the number of heavy atoms, and R is the number of recycling iterations. Here, *y_ij_* and *d_ijr_* denote the distances between atoms *i* and *j* in the experimental structure and in the structure predicted after recycling iteration *r*, respectively. The overlap scaling factor was set to 0.6. The bond loss constrained intranucleotide bond lengths to remain close to their experimentally observed values: 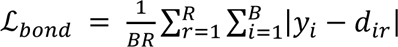, where B is the number of covalent bonds in the structure, and *y_i_* and *d_ir_* denote the length of bond *i* in the experimental structure and in the structure predicted after recycling iteration *r*, respectively. The pLDDT loss, originally introduced in AlphaFold2, was used to supervise the confidence-prediction module. FAPE was also introduced in AlphaFold2 to constrain predicted structures relative to their corresponding experimental structures. We replaced the original FAPE loss with a weighted FAPE loss (Supplementary Algorithm 4). Whereas the original all-atom FAPE loss assigns equal weights to atom pairs within the cutoff distance, the weighted FAPE loss assigns greater weights to smaller structural deviations, thereby emphasizing local geometric refinement and the reduction of steric clashes. To further improve structural accuracy, the dRMSD loss, originally introduced in RGN^58^ for protein structure prediction, was computed from pairwise distances between C3′ atoms to promote accurate local geometry and facilitate optimization. The weights assigned to the clash, bond, pLDDT, weighted FAPE, and dRMSD losses were 0.01, 0.01, 0.01, 1.0, and 0.001, respectively. The resulting total loss for each training sample was further multiplied by the square root of the cropped sequence length.

### Evaluation metrics

The accuracy of the predicted structures was evaluated using TM-score, RMSD, MCQ, and INF. TM-score and RMSD between the predicted structures and the corresponding experimental structures were computed using USAlign (https://anaconda.org/bioconda/usalign)^59^. MCQ was computed using functions provided in the RNA-TorsionBERT^50^ Python scripts (https://github.com/EvryRNA/RNA-TorsionBERT/blob/main/src/metrics/mcq.py), and INF was computed using the RNA_assessment^48^ package (https://github.com/RNA-Puzzles/RNA_assessment).

### Benchmark methods

DRFold was obtained from its GitHub repository (https://github.com/leeyang/DRfold/tree/main). DRFold2 was obtained from its GitHub repository (https://github.com/leeyang/DRfold2). NUFold was obtained from its GitHub repository (https://github.com/kiharalab/nufold/). trRosettaRNA was obtained from the trRosettaRNA web server (https://yanglab.qd.sdu.edu.cn/trRosettaRNA/). AlphaFold3 was obtained from its GitHub repository (https://github.com/google-deepmind/alphafold3). RhoFold+ was obtained from its GitHub repository (https://github.com/ml4bio/RhoFold). RoseTTAFoldNA was obtained from its GitHub repository (https://github.com/uw-ipd/RoseTTAFold2NA/tree/main).

## Acknowledgements

We thank Yiduo Xiong at Huazhong University of Science and Technology for insightful discussion. We thank the developers and contributors of open-source tools and datasets including the PDB, PyTorch, and others for greatly accelerating our research. This work was supported by National Key R&D Program of China (2022YFA1004800, 2025YFF1207900), Natural Science Foundation of China (T2350003, T2341007, 12131020, 42450084, 42450135, 12326614, 12404236 and 12426310), Zhejiang Province Vanguard Goose-Leading Initiative (2025C01114), Science and Technology Commission of Shanghai Municipality (23JS1401300), and Hangzhou Institute for advanced study of UCAS (2024HIAS-P004, 2024ZZBS06), and JST Moonshot R&D (JPMJMS2021).

## Author contributions

L.C. and Y.S. conceived the study. Y.S. performed the experiments, analyzed the data and wrote the original draft. Y.S. and S.Z. reviewed and edited the manuscript. L.C. and S.Z. contributed to data visualization. L.C. supervised the study and secured funding.

## Competing interests

The authors declare no competing interests.

## Data and code availability

The source code and pretrained weights for NUMonomer are publicly available at https://github.com/yunda-si/NUMonomer.

## Supplemental Information

### Figures

**Figure S1:**
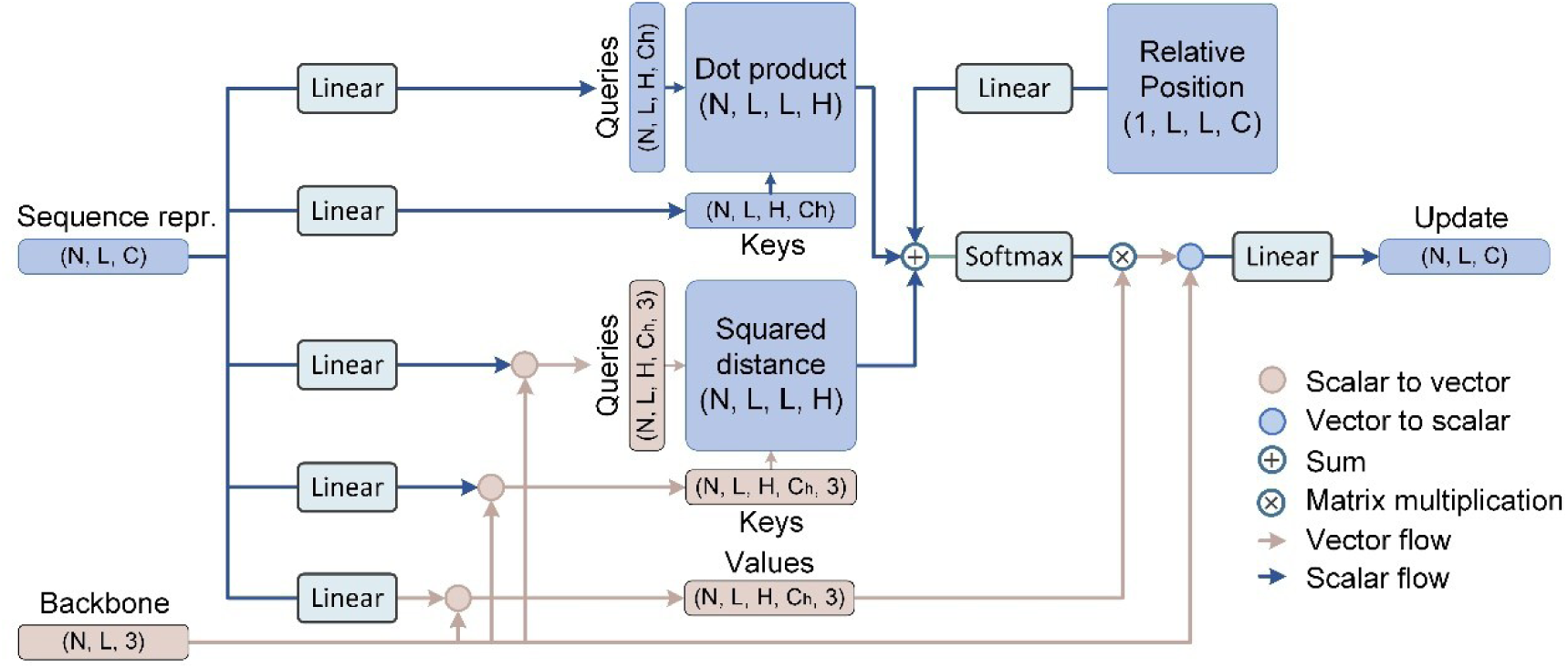
Architecture of the Inter-nucleotide attention module. The module takes sequence representation and coordinates as input and updates the sequence representation via learned inter-nucleotide interactions. Here, N denotes the number of atoms, L the sequence length, C the number of channels (default 384), and H the number of attention heads (default 12).

**Figure S2:**
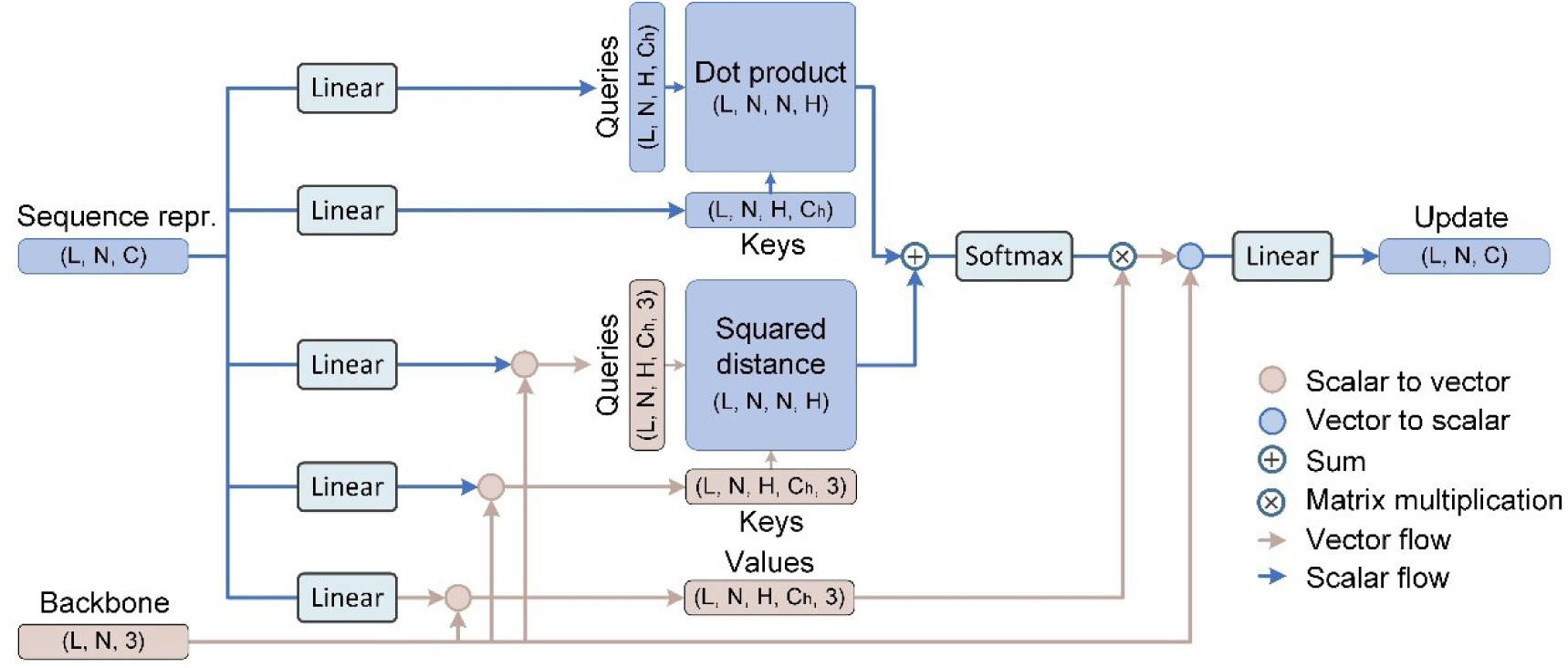
Architecture of the Intra-nucleotide attention module. The module takes sequence representation and coordinates as input and updates the sequence representation via learned intra-nucleotide interactions. Here, N denotes the number of atoms, L the sequence length, C the number of channels (default 384), and H the number of attention heads (default 12).

**Figure S3:**
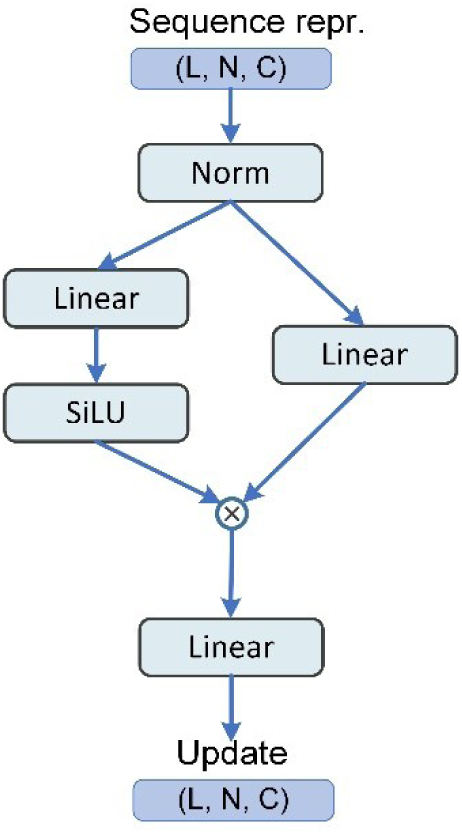
Architecture of the transition module. The module takes sequence representation as input and updates the sequence representation along the channel dimension.

**Figure S4:**
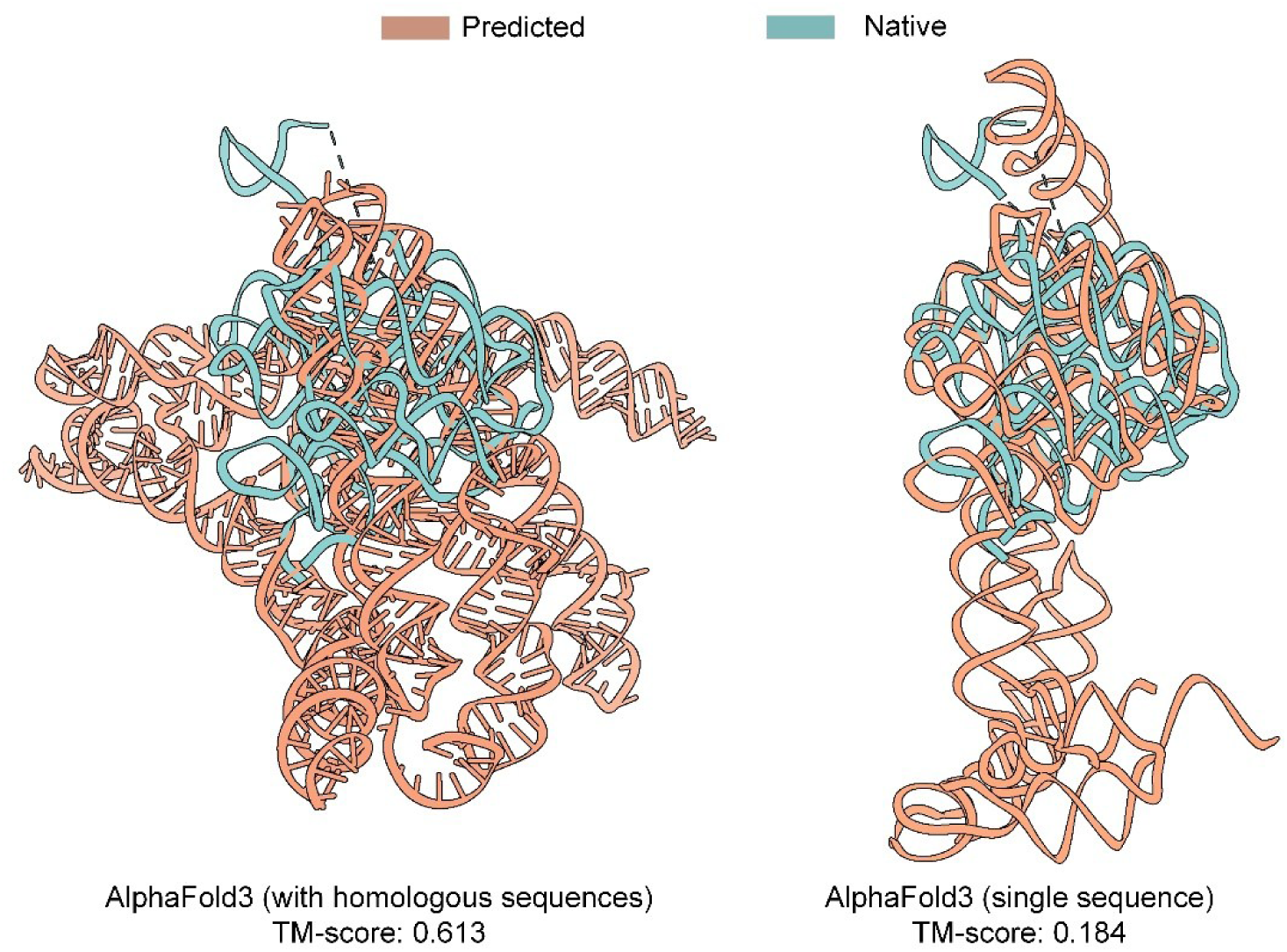
AlphaFold3 predictions for CASP16 target R1285. R1285 is 577 nucleotides long. The prediction generated with an MSA achieved a TM-score of 0.613, whereas the prediction generated from the query sequence alone achieved a TM-score of 0.184.

**Figure S5:**
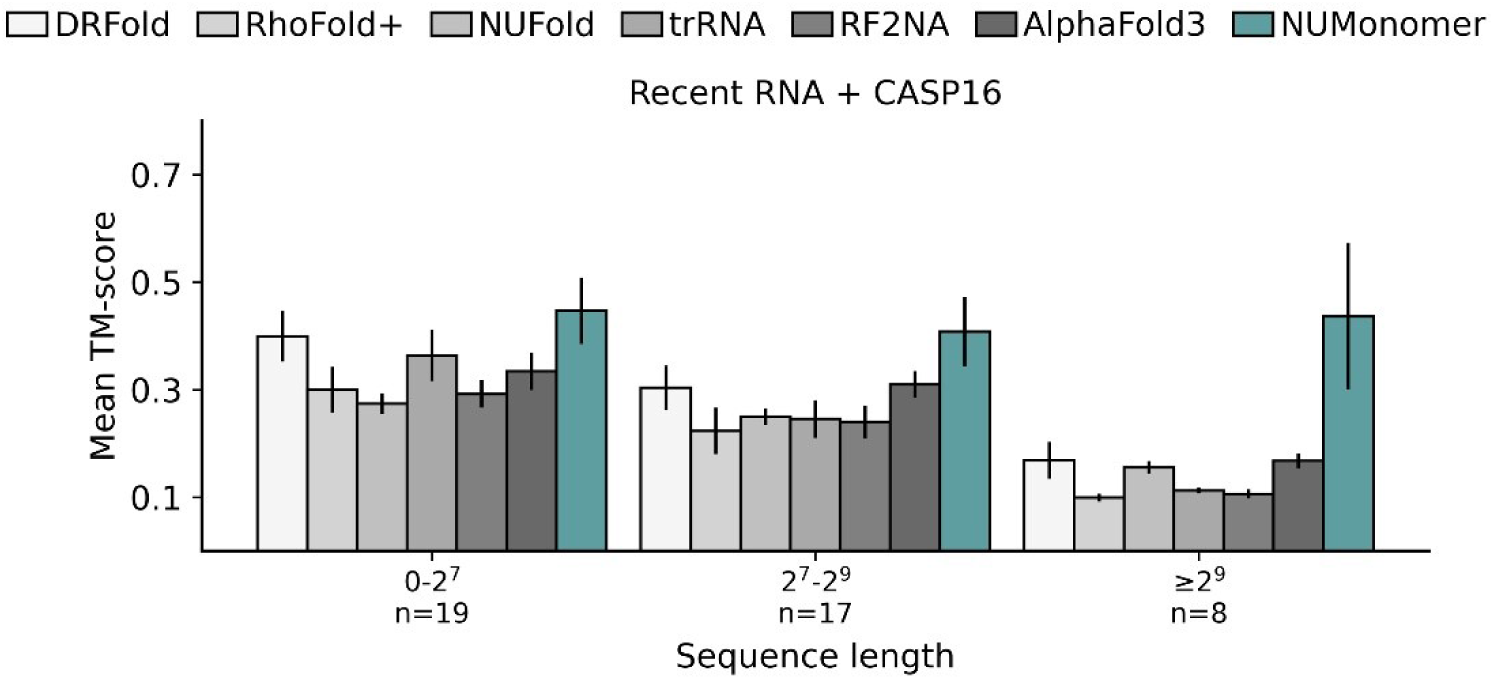
Mean TM-scores of NUMonomer, NUFold, RhoFold+, AlphaFold3, trRosettaRNA, DRFold, and RoseTTAFoldNA under single-sequence inference across RNA sequence-length groups in the combined Recent RNA and CASP16 datasets. Black vertical lines indicate the standard deviation across RNA targets. n values indicate the number of RNA structures in each interval.

**Figure S6:**
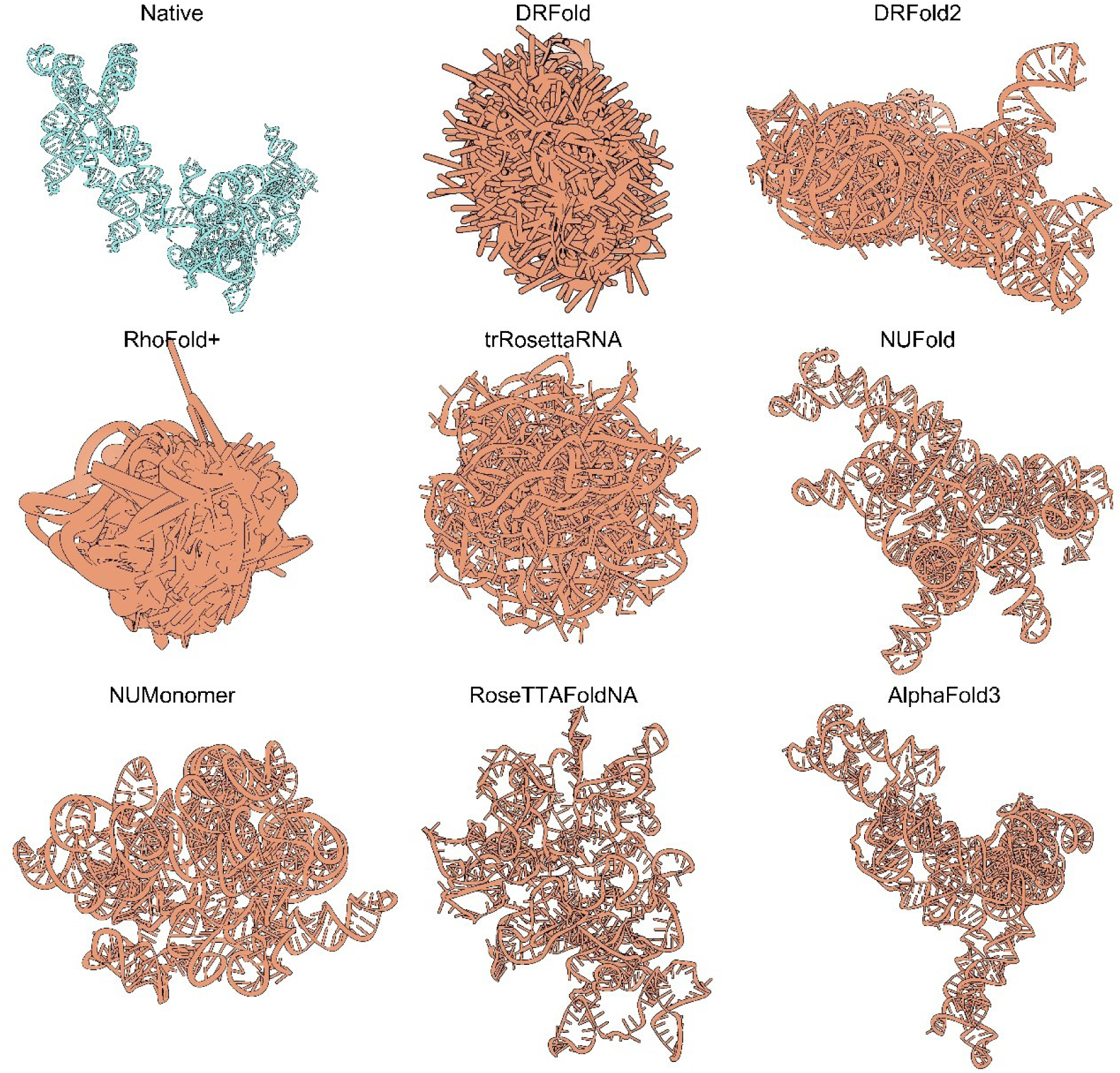
Examples of failed predictions for CASP16 target R1251. Under their default inference settings, all evaluated methods failed to recover the global fold of CASP16 target R1251, yielding TM-scores ranging from 0.00 to 0.19. Nevertheless, predictions generated by NUMonomer, NUFold, RoseTTAFoldNA, and AlphaFold3 still captured local base-pairing interactions, resulting in the formation of plausible helical segments despite the incorrect global topology.

**Figure S7:**
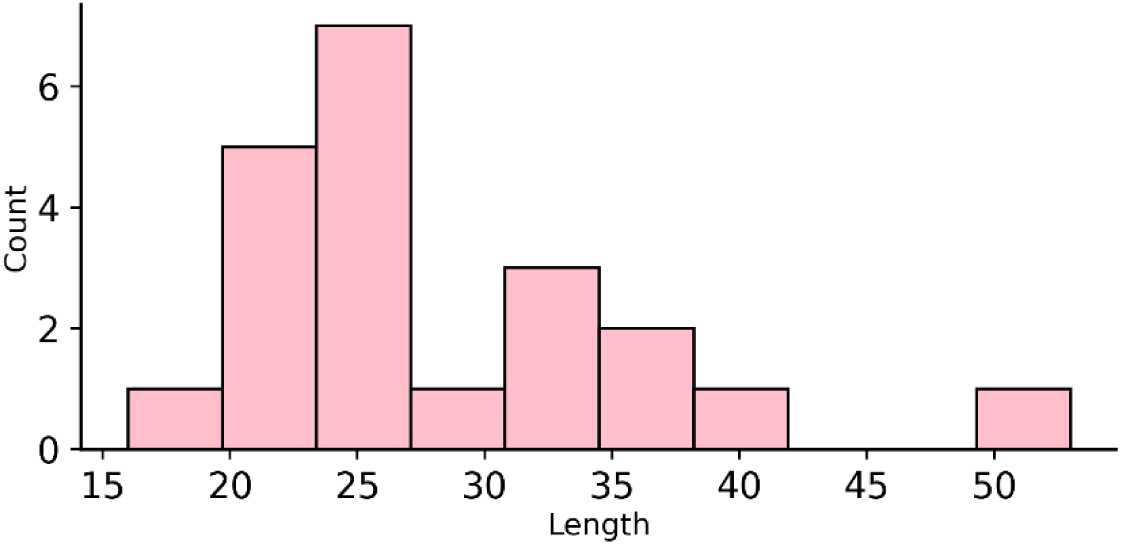
Length distribution of ssDNA sequences in the Recent ssDNA dataset.

**Figure S8:**
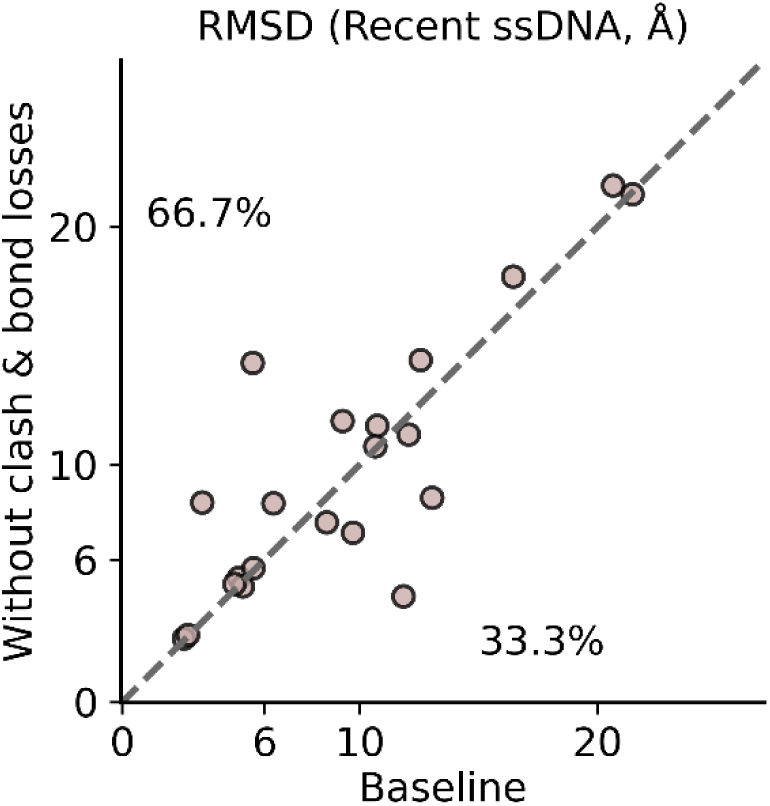
Head-to-head comparison of RMSD values between the baseline model and the model trained without clash and bond losses on the Recent ssDNA dataset. Percentages indicate the proportion of targets in the upper and lower triangular regions.

**Figure S9:**
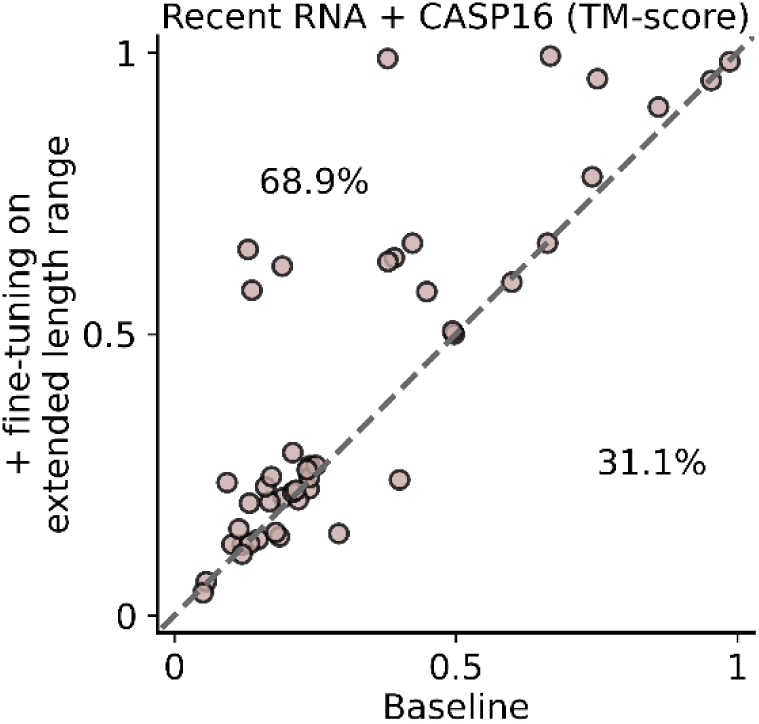
Head-to-head comparison of TM-scores between the baseline model and the model fine-tuned on an extended sequence-length range on the combined Recent RNA and CASP16 datasets. Percentages indicate the proportion of targets in the upper and lower triangular regions.

**Figure S10:**
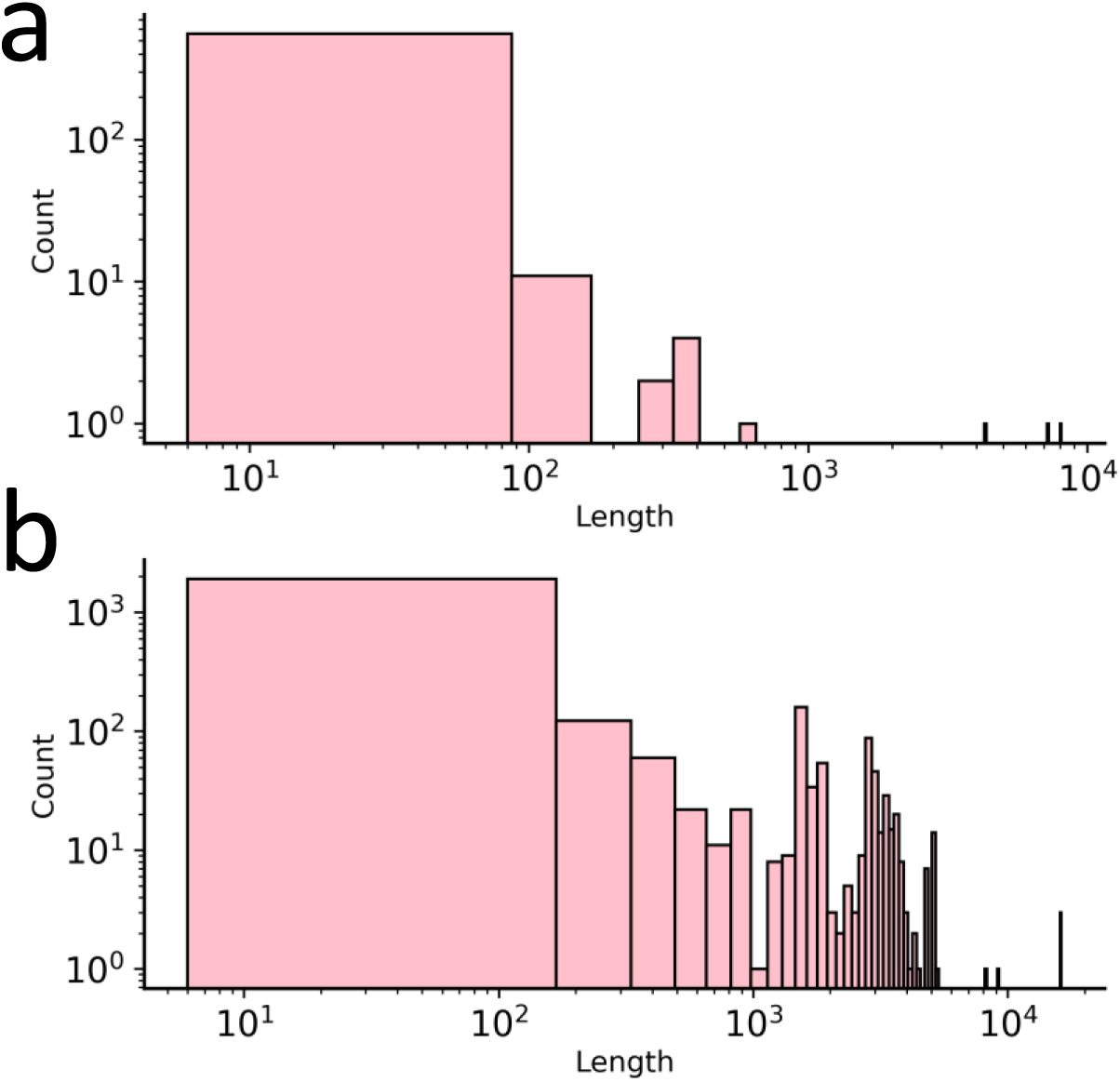
Length distribution of RNA (a) and ssDNA (b) sequences in the NUMonomer training dataset.

**Figure S11:**
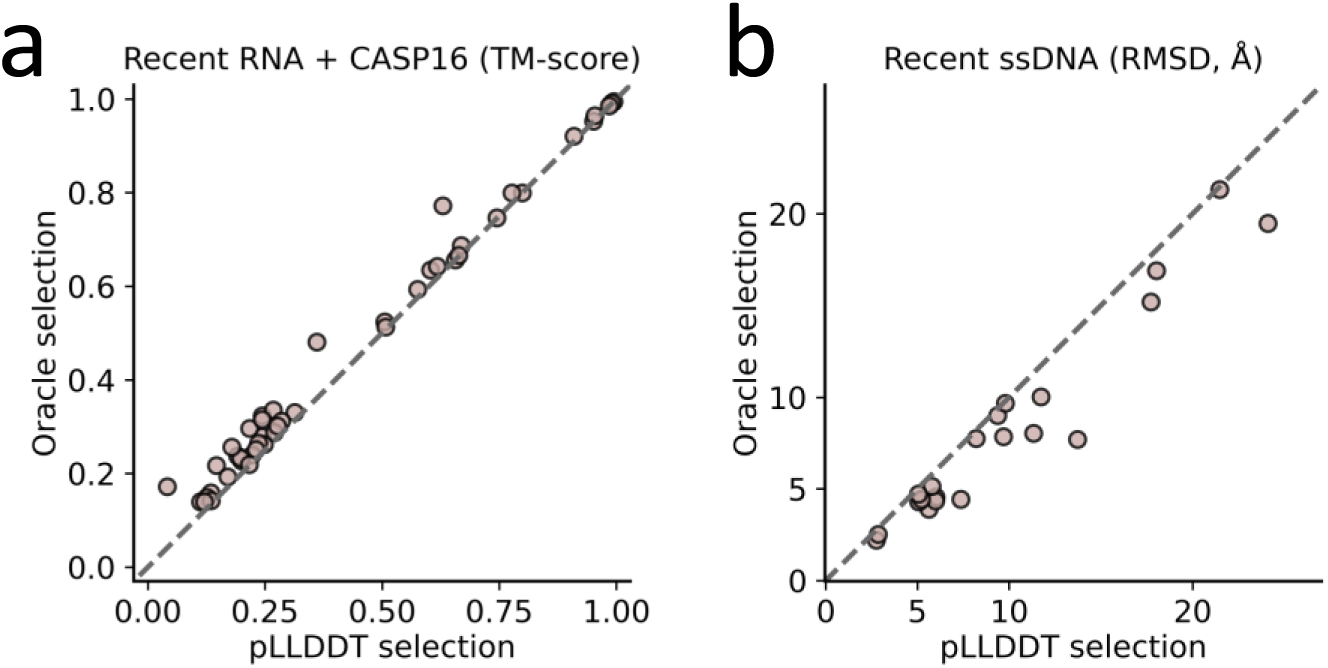
Head-to-head comparison of the performance of the oracle selection strategy and the predicted confidence score-based selection strategy on the combined Recent RNA and CASP16 datasets (**a**) and the Recent ssDNA dataset (**b**).

### Tables

**Table S1:**
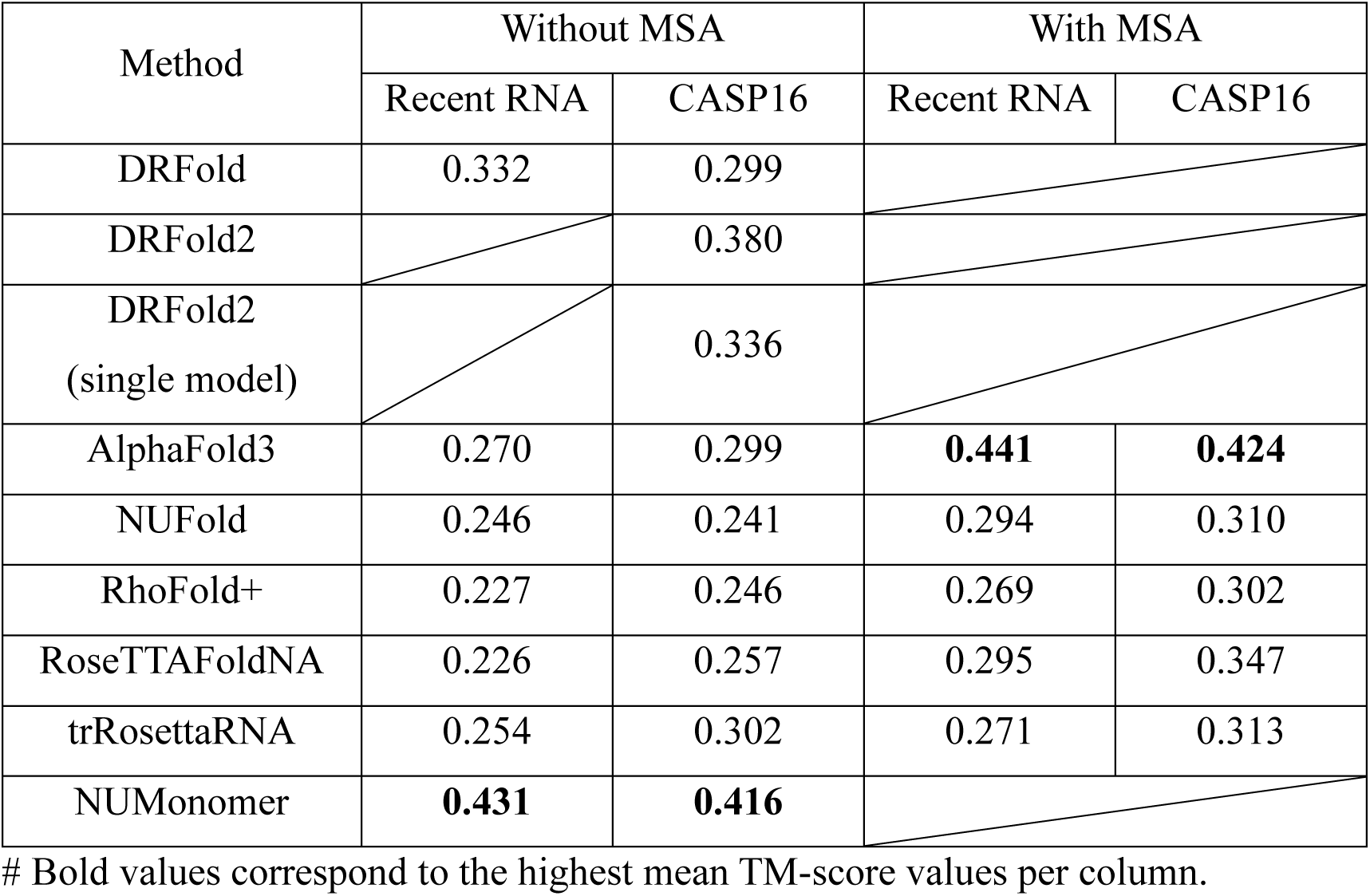
Mean TM-score of NUMonomer and the other evaluated methods on the Recent RNA and CASP16 datasets.

**Table S2:** Mean TM-scores of NUMonomer, NUFold, RhoFold+, AlphaFold3, trRosettaRNA, DRFold, and RoseTTAFoldNA across RNA sequence-length groups in the combined Recent RNA and CASP16 datasets under their standard inference settings.

| Method | Short group | Medium group | Long group |
| --- | --- | --- | --- |
| DRFold | 0.399 | 0.303 | 0.169 |
| AlphaFold3 | 0.393 | **0.444** | **0.517** |
| NUFold | 0.371 | 0.270 | 0.183 |
| RhoFold+ | 0.352 | 0.283 | 0.097 |
| RoseTTAFoldNA | 0.393 | 0.311 | 0.134 |
| trRosettaRNA | 0.389 | 0.250 | 0.110 |
| NUMonomer | **0.447** | 0.407 | 0.437 |

**Table S3:** Inference times of NUMonomer, NUFold, AlphaFold3, and RoseTTAFoldNA under single-sequence inference. Values are reported in seconds.

| Length (nt) | NUMonomer | NUFold | AlphaFold3 | RoseTTAFoldNA |
| --- | --- | --- | --- | --- |
| 100 | 1.1 | 62.84 | 56.16 | 72.32 |
| 500 | 2.05 | 188.49 | 67.12 | 152.84 |
| 1000 | 3.79 | 486.96 | 150.21 | 299.81 |
| 2000 | 4.28 | 1894.5 | 234.07 | 871.27 |
| 5000 | 18.84 |  |  |  |

**Table S4:** Mean and median MCQ of NUMonomer and the other evaluated methods on the combined Recent RNA and CASP16 datasets.

| Method | Without MSA |  | With MSA |  |
| --- | --- | --- | --- | --- |
|  | Median | Mean | Median | Mean |
| DRFold | 30.0 | 33.7 |  |  |
| AlphaFold3 | 18.0 | 19.7 | **17.4** | **18.8** |
| NUFold | 20.4 | 22.8 | 19.8 | 22.0 |
| RhoFold+ | 71.6 | 77.7 | 65.7 | 74.2 |
| RoseTTAFoldNA | 20.5 | 23.3 | 20.5 | 23.5 |
| trRosettaRNA | 35.3 | 48.5 | 33.8 | 47.9 |
| NUMonomer | **17.1** | **19.5** |  |  |

**Table S5:** Mean and median INF of NUMonomer and the other evaluated methods on the combined Recent RNA and CASP16 datasets.

| Method | Without MSA |  | With MSA |  |
| --- | --- | --- | --- | --- |
|  | Median | Mean | Median | Mean |
| DRFold | 0.618 | 0.48 |  |  |
| AlphaFold3 | **0.721** | **0.672** | **0.801** | **0.749** |
| NUFold | 0.633 | 0.566 | 0.692 | 0.614 |
| RhoFold+ | 0.329 | 0.326 | 0.560 | 0.429 |
| RoseTTAFoldNA | 0.575 | 0.549 | 0.643 | 0.577 |
| trRosettaRNA | 0.543 | 0.386 | 0.573 | 0.417 |
| NUMonomer | 0.686 | 0.660 |  |  |

**Table S6:** Performance of NUMonomer and AlphaFold3 on the Recent ssDNA dataset.

|  | Median |  | Mean |  |
| --- | --- | --- | --- | --- |
|  | AlphaFold3 | NUMonomer | AlphaFold3 | NUMonomer |
| RMSD | 7.47 | 8.20 | 8.01 | 9.85 |
| MCQ | 23.5 | 27.7 | 25.4 | 28.1 |
| INF | 0.593 | 0.557 | 0.560 | 0.505 |

**Table S7:** Performance of NUMonomer ablation variants on the RNA and ssDNA benchmark datasets.

|  | RNA (TM-score) |  | ssDNA (RMSD) |  |
| --- | --- | --- | --- | --- |
|  | Median | Mean | Median | Mean |
| - clash & bond loss | 0.171 | 0.301 | 8.41 | 9.81 |
| Baseline | 0.220 | 0.324 | 9.27 | 9.42 |
| + dynamic length | 0.232 | 0.367 | **8.14** | **9.22** |
| + fine-tuning on extended length range | 0.247 | 0.406 | 8.29 | 9.44 |
| + sampling during inference | **0.270** | **0.425** | 8.20 | 9.85 |

### Model architecture

#### Main Framework

##### Algorithm 1 Inference

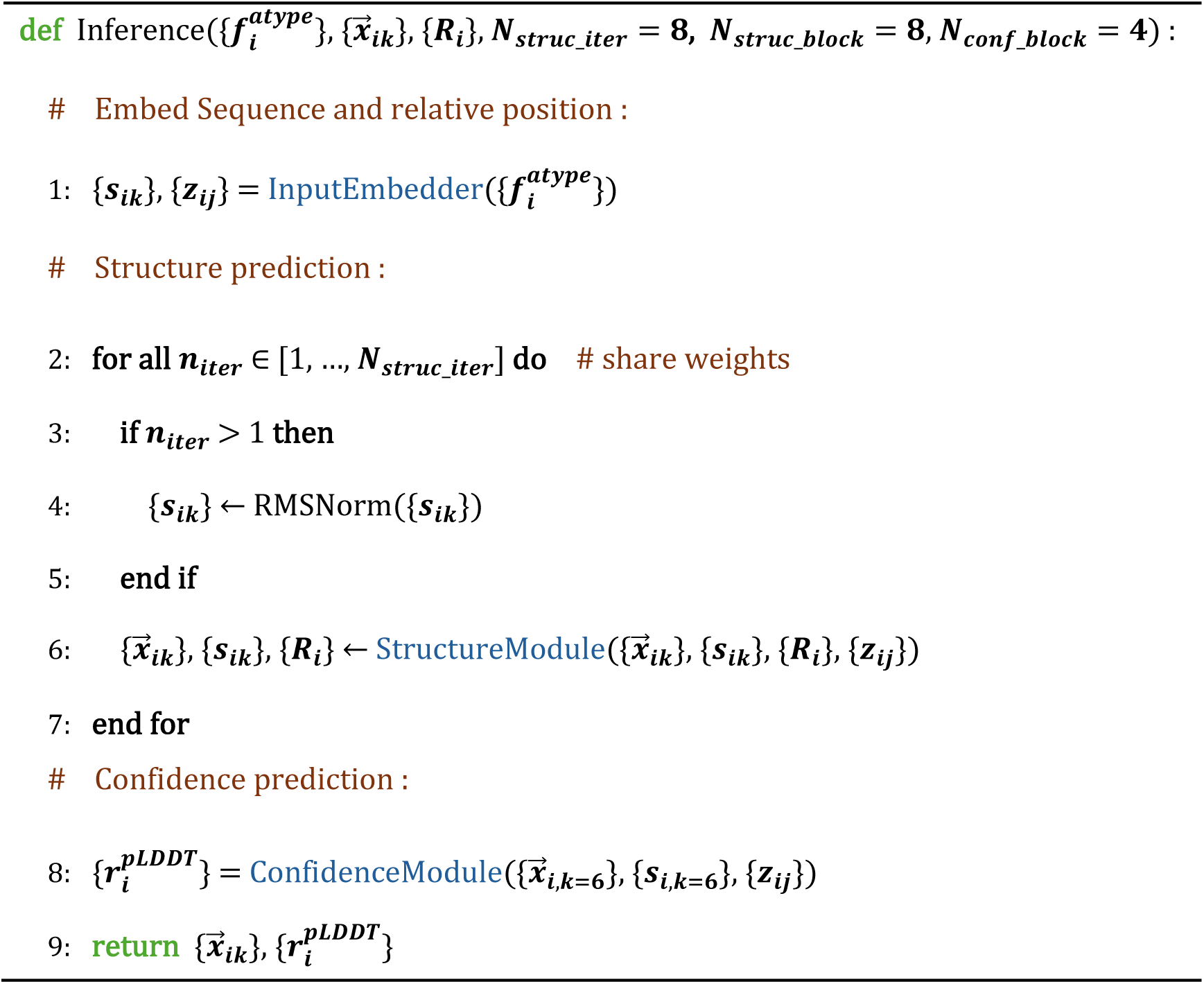

NUMonomer takes a single nucleic acid sequence together with randomly initialized all-atom coordinates as input and predicts all-atom coordinates and per-nucleotide pLDDT confidence scores. The InputEmbedder encodes the input sequence and relative positional information into pairwise and single representations. The StructureModule iteratively updates the sequence representations to infer nucleotide interactions and predicts all-atom coordinates. The ConfidenceModule estimates per-nucleotide confidence scores for the predicted structures and the corresponding structure representations generated by the StructureModule, with stop-gradient applied. Throughout this work, 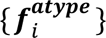 denote the nucleotide type at sequence position 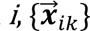 denote the coordinates of atom *k* in nucleotide *i*, ***R****_i_* denotes the local coordinate frame of nucleotide *i*, which is constructed using the Gram–Schmidt procedure from the coordinates of the P, C3′, and C1′ atoms, with the C3′ atom serving as the origin, and ***z****_ij_* denotes the relative positional encoding between nucleotides *i* and *j*, following the positional encoding scheme introduced in AlphaFold2. Unless otherwise specified, atom index *k* = 6 corresponds to the C3′ atom.

#### StructureModule

##### Algorithm 2 StructureModule

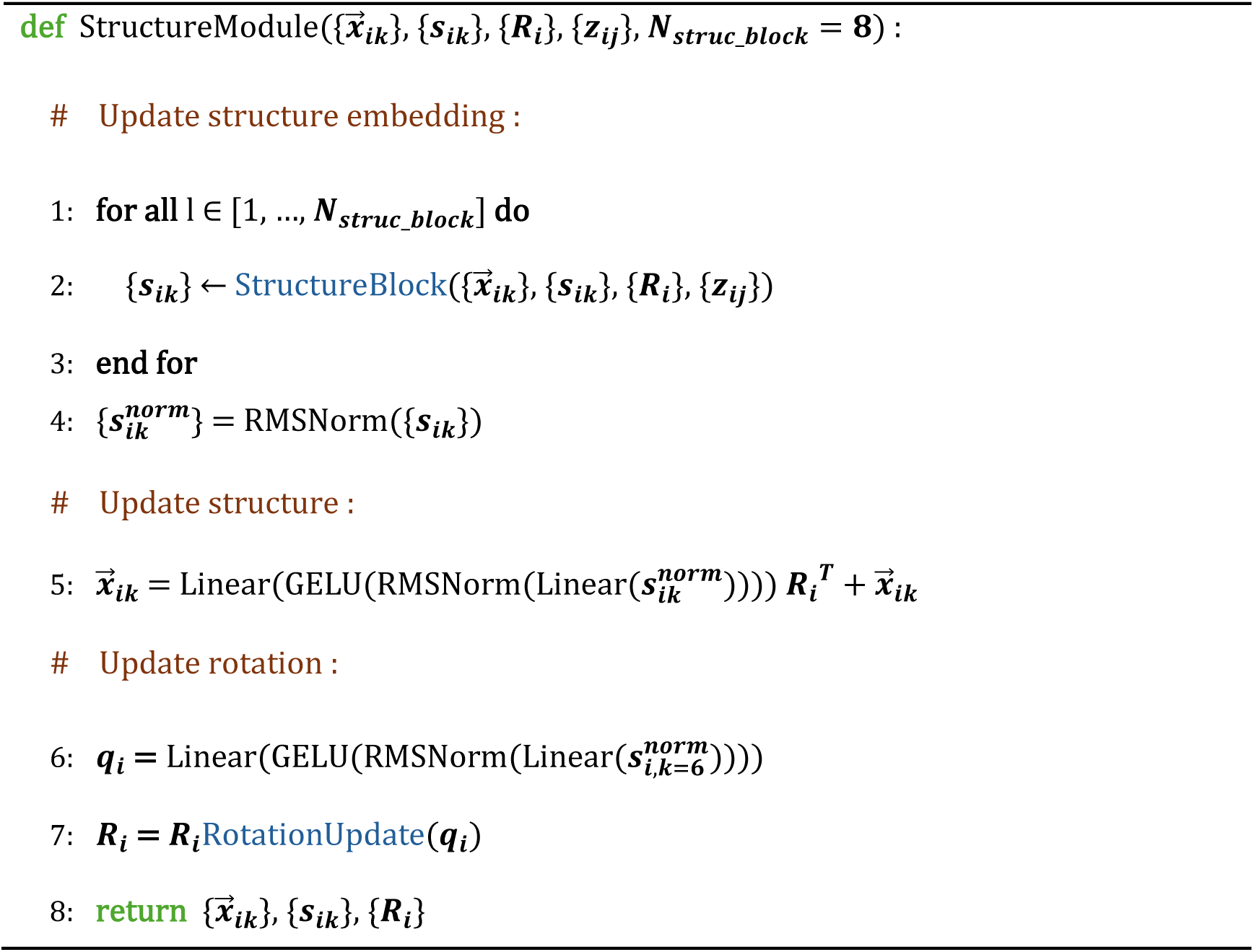

The StructureModule consists of eight structure blocks that iteratively refine the structure representations and predict the all-atom structure.

##### Algorithm 3 Rotation update

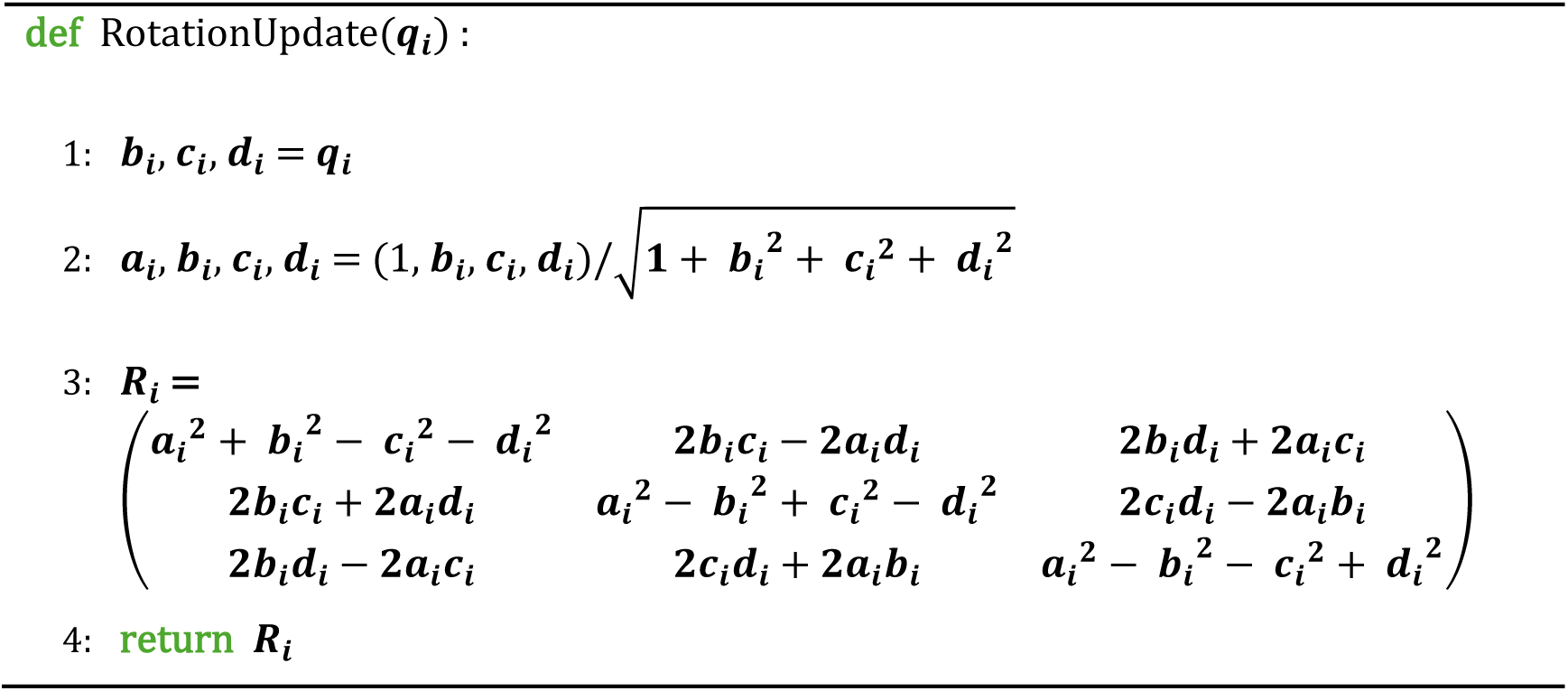

#### Weighted FAPE Loss

##### Algorithm 4 Weighted FAPE Loss

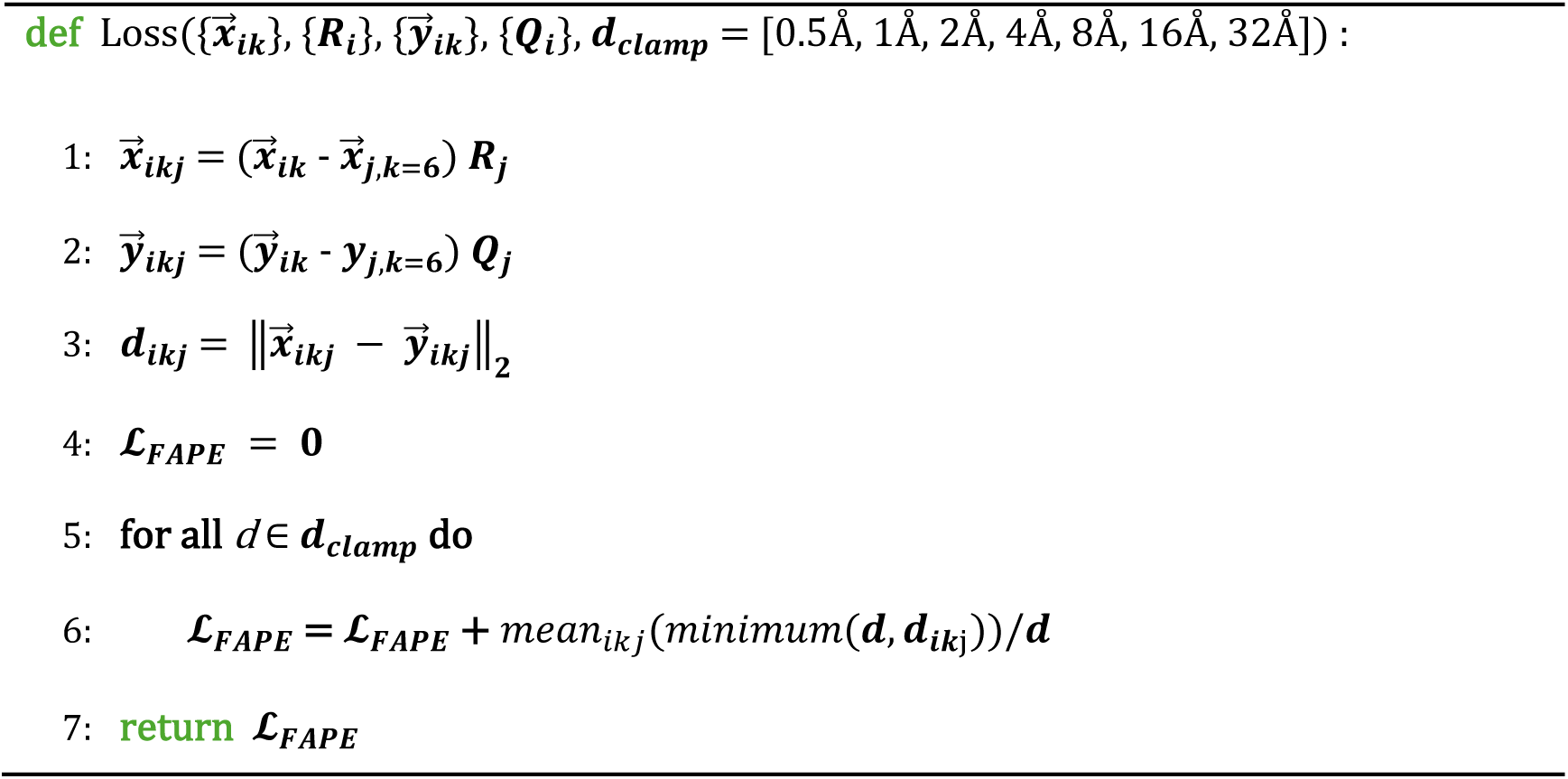

Here, 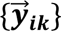 denotes the corresponding coordinates in the experimental structure, and **Q***_i_* denotes the corresponding local coordinate frame.

## Notes

https://github.com/yunda-si/NUMonomer

### Competing Interest Statement

The authors have declared no competing interest.

